# Conditional Localization Pharmacology Manipulates Cell Cycle with Spatiotemporal Precision

**DOI:** 10.1101/2024.09.12.612697

**Authors:** Changfeng Deng, Yung-Chi Lan, Geng-Yuan Chen, Chigozie S. Ekeabu, Quoc Doan, Megan Chung, Micheal A. Lampson, David M. Chenoweth

## Abstract

Traditional pharmacology has limited control of drug activity and localization in space and time. Herein, we described an approach for kinase regulation using conditional localization pharmacology (CLP), where an inactive caged inhibitor is localized to a site of interest in a dormant state using intracellular protein tethering. The activity of the inhibitor can be regulated with spatial and temporal precision in a live cellular environment using light. As a proof of concept, a photocaged MPS1 kinase inhibitor (reversine) bearing a Halo-tag ligand tether was designed to manipulate the cell cycle. We demonstrate that this new caged reversine halo probe (CRH) strategy is capable of efficient localization and exceptional spatiotemporal control over spindle assembly checkpoint (SAC) silencing and mitotic exit.

**Table of Contents artwork:** 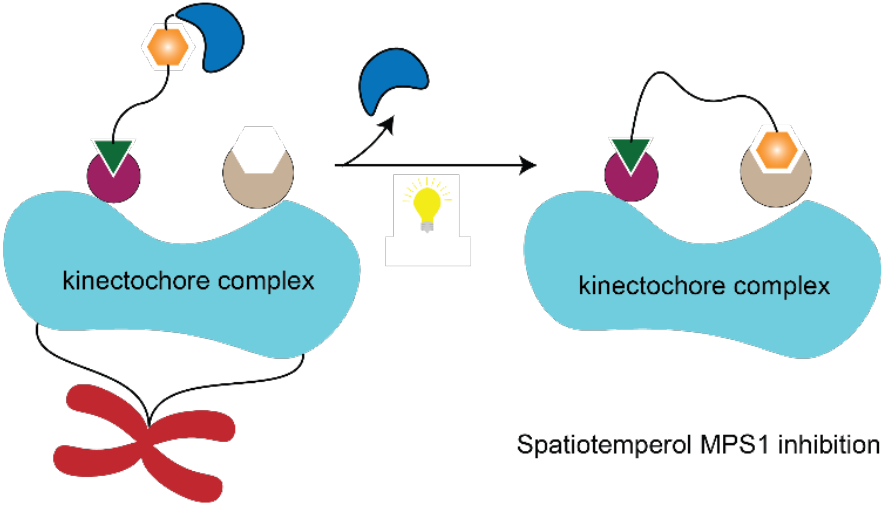

## INTRODUCTION

Manipulation of protein behavior in space and time is critical to understanding biological processes. Small molecule modulators are widely used to control protein function and dissect intricate biological pathways. Traditional chemical or pharmacological approaches to modulate protein function involve direct treatment with drug molecules. These molecules can freely diffuse and exchange between cells and culture medium through cell uptake/efflux with little control over subcellular localization (**Figure 1a**). A wide range of proteins share high structure and sequence similarity, and this lack of spatial and temporal control over their ligands can severely diminish on target potency and selectivity, often confounding experimental results.^1^

**Figure 1.**
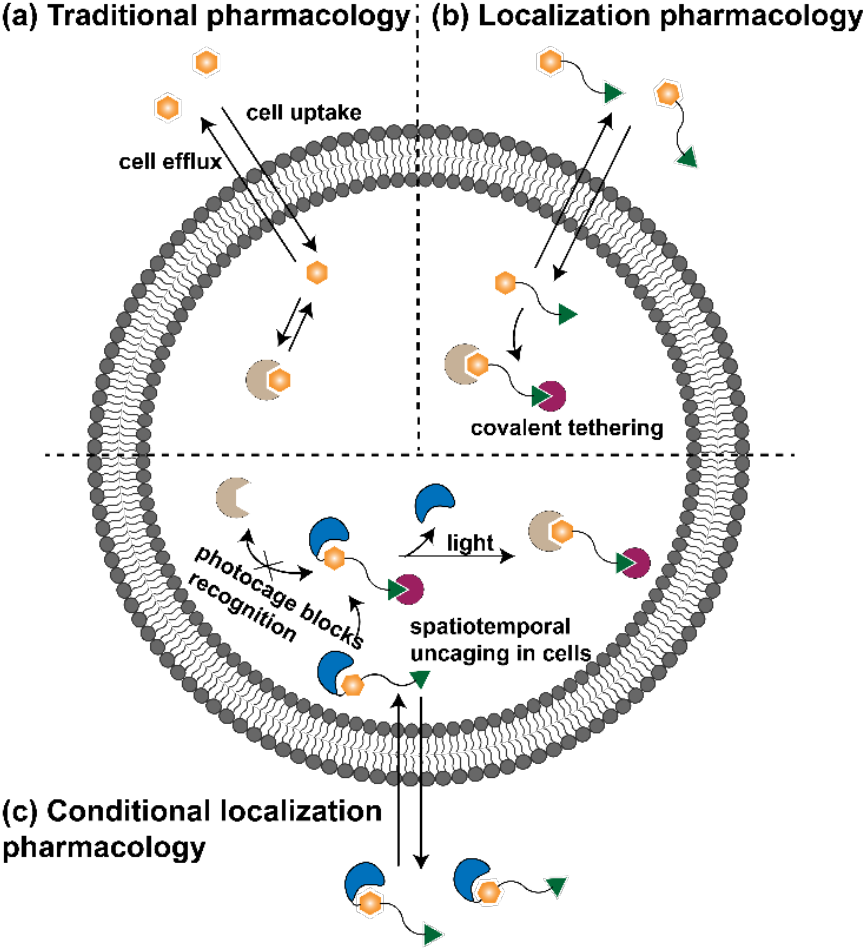
Schematics of pharmacological strategies. (a) Traditional pharmacology relies on inhibitors that freely diffuse in and out of cells. (b) Localization pharmacology spatially constrains the inhibitor by tethering to a protein anchor, which prevents efflux. (c) Conditional localization pharmacology links a photocage to the inhibitor moiety of the tethered inhibitor, so that inhibition can be controlled by a light pulse. when incorporating Halo-tag localization.^5^

Localization pharmacology has been developed as a powerful chemogenetic tool to address challenges associated with the spatial distribution and targeted activity of pharmacological agents^2^. Using localization pharmacology, small molecule modulators containing a reactive tether can attach covalently or noncovalently to a target protein^1^. Fusion of a self-labeling (tag) protein such as SNAP-tag^3^, CLIP-tag^4^ or Halo-tag^5^ to the target protein of interest is often used in covalent tethering strategies. The tag proteins exhibit fast binding kinetics and superior selectivity toward the tethered small molecule modulator, enabling fast and effective localization at low concentrations. For example, Halo-tag tethering can generate a high spatial concentration of a small molecule drug via proximity, and the working concentration of a compound with modest potency (50% inhibitory concentration IC_50_ > 1 uM) can be reduced to 100 nM.

As shown in **Figure 1b**, once a tethered small molecule modulator enters a cell containing the complement tagging protein, covalent immobilization ensues, sequestering the small molecule inside the cell and preventing efflux. One drawback of the tagging strategy in protein-based localization pharmacology is that the immobilized pharmacological agent remains active, whereas spatial and temporal control of activity is highly desired for many experiments.

To complement the lack of pharmacological activity control in localization pharmacology, photoswitchable pharmacological agents have been developed for light-responsive turn-on and turn-off control. Pioneering efforts by the Trauner group^7, 8, 9^ have advanced the field of photopharmacology and tethered photopharmacology by developing a wide array of membrane protein optogenetic tools. These tools have found widespread use in neuropharmacology, and many elegant photoswitch-functionalized ligand designs have emerged for control of cell surface receptors.

As a complementary technique to the tethered photoswitch approach, we have developed a caged-ligand tethering approach for discrete control of intracellular protein activity. In some cases, tight experimental control over activity of the tethered pharmacological agent is required, and utilization of an appropriate caging strategy can produce a stable and completely inactive state, a requirement that is hard to achieve with other approaches. To provide a clean off/on conditional tethered photopharmacological effect, we combine a photocaged protein ligand with a localization pharmacology (tethering) approach. Our tethered caged-ligand design consists of three components as follows: a photocage (head), a drug (body), and a Halo-tag ligand (reactive tether). As illustrated in **Figure 1c**, the inactive probe molecules can cross the membrane and are bound inside the cell through covalent immobilization to the tagging protein. A wash-out step can be used to remove any excess unbound probe, or low-level dosing can be employed to ensure no excess probe remains. This leads to a caged (inactive) probe molecule tethered at an intracellular location of interest. A photoactivation step can then be used to activate the localized drug from its dormant (caged) state in a spatially defined and temporally controlled manner. The spatiotemporal control afforded by this technique allows for precise control of activity at the single cell level or within a discrete population of cells without affecting any neighboring cells. After uncaging, the tethered activated pharmacological agent is free to bind the recognition pocket of the protein of interest (POI), leading to superior spatially confined and temporally precise modulation of the protein function compared to photo-triggered drug release alone^10,11^. As a proof of principle, we have developed a new tool for manipulation of cell division using caged localization pharmacology (CLP).

During cell division, the spindle assembly checkpoint (SAC) is activated until all chromosomes are properly attached to the spindle, the microtubule-based machinery that physically segregates the chromosomes. After chromosomes align in the middle of the spindle at metaphase, the SAC is silenced to allow progression to anaphase and separation of sister chromosomes to opposite sides of the cell, which will ultimately become two daughter cells. Events before and after anaphase rely on many of the same players, including microtubule-associated proteins and kinetochore proteins that mediate attachment of chromosomes to the spindle. Experimental perturbations of these proteins often activate the SAC, so that the cell arrests in metaphase and does not enter anaphase. To study anaphase under these conditions, the SAC can be silenced by inhibiting MPS1, a key checkpoint kinase that localizes to kinetochores to activate the SAC. However, diffusion-based MPS1 inhibitors such reversine^12^ lack spatiotemporal precision that pose challenges for experimental efficiency. Once the inhibitor is introduced to the cells, all cells enter anaphase at the same time. Anaphase is highly dynamic and lasts only a few minutes, so experiments typically require live imaging of individual cells with high time resolution. If all cells enter anaphase simultaneously, only one cell can be imaged in each experiment. One potential solution to this difficulty involves releasing cells from metaphase arrest one at a time using CLP.

## RESULTS AND DISCUSSION

We designed a mitotic regulatory tool named caged reversine-halo (CRH) consisting of a halo ligand that binds to a halo-tagged protein, reversine^12^ (structure shown in **Figure 2b**) and a photocage to realize light-controlled release. Examining the crystal structure of the MPS1 and reversine complex reveals that the purine ring is located in the binding pocket, resulting in electrostatic interactions with the backbond of amino acids 603E and 605G. Herein, the core purine ring is in an ideal position to be shielded by a photocage, which will effectively block recognition through disruption of key H-bonds. For photocaging on the purine ring side, a carbamate photocage linker is not suitable due to stability issues, and therefore, we adopt the similar direct nitrogen caging strategy^13^ over benzylguanine in the SNAPTag with 6-nitro-1,3-benzodioxole. The exposed morpholine ring represents the ideal site for tethering and can be replaced with a piperazine ring^14^ for chemical modification without losing reversine activity. We designed a six-peg unit linker and a two-peg unit linker tethered chlorohexyl warhead at the piperazine nitrogen.

**Figure 2.**
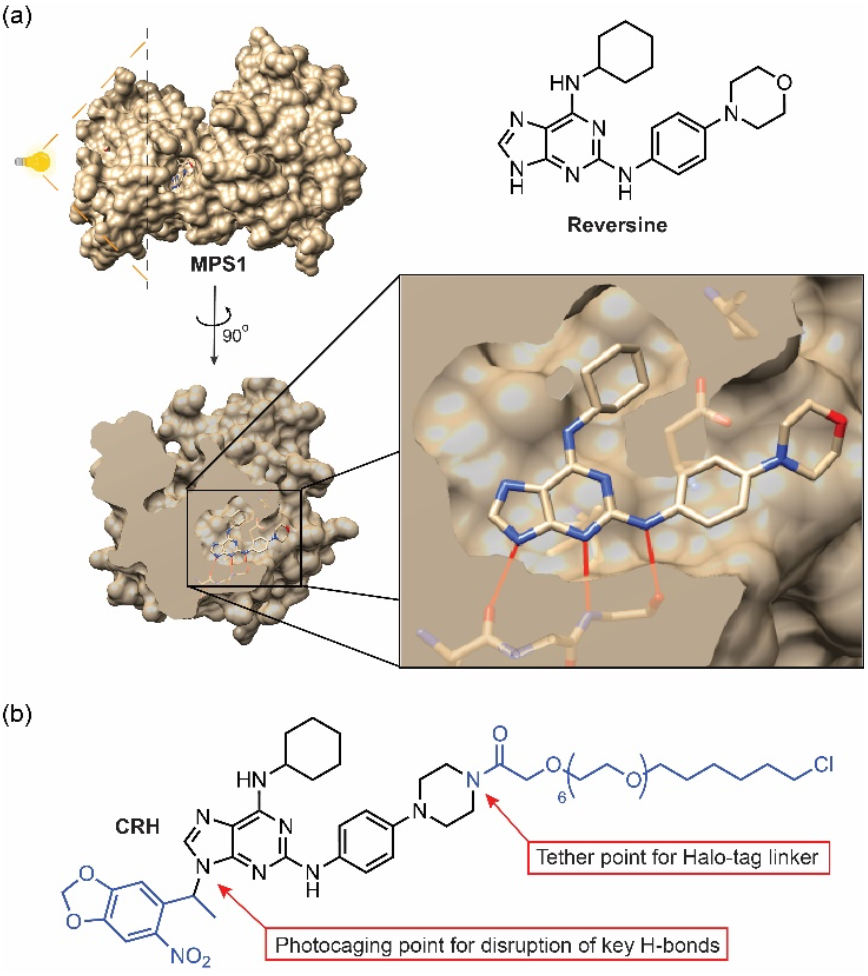
Molecular design of the photocaged MPS1 kinase inhibitor (CRH). (a) Structure of the MPS1 inhibitor reversine and crystal structure of reversine binding to the MPS1 active site (PDB, 5ljj). Cutaway view shows critical hydrogen bonding interactions of MPS1 with reversine and exit tunnel from reversine binding site. (b) Structure of CRH: reversine tethered to a Halo-tag ligand containing a PEG linkers at piperazine moiety as well as a photocage at benzylguanine moiety that blocks key ligand binding interactions in the MPS1 active site shown in (a).

We first tested if the photocage shields reversine activity. We used an inhibitor of a mitotic kinesin, CENP-E (kinesin-7), to arrest cells in mitosis with misaligned chromosomes (**Figure 3a**). Reversine silences the SAC and promotes anaphase onset and mitotic exit, reducing mitotic cells in the population (measured as mitotic index). Thus, cells treated with reversine or uncaged reversine-halo (RH) should have a lower mitotic index than control cells, while CRH should have no effect. Addition of either reversine or RH reduced the mitotic index from ~16% to ~2%, indicating checkpoint silencing. In contrast, the mitotic index remained high in cells treated with CRH (**Figure 3b**). These results indicate that the reversine moiety preserves its SAC silencing function after the tethering modification and that the cage successfully shields reversine from MPS1. For these experiments we used a Hela cell line expressing a kinetochore protein, SPC25, fused to HaloTag. Covalent anchoring of CRH or RH to SPC25, which sits in proximity to MPS1 binding sites, provides the spatial precision of MPS1 inhibition. This anchoring is thermodynamically irreversible, thereby resisting efflux and confining CRH within the cell.

**Figure 3.**
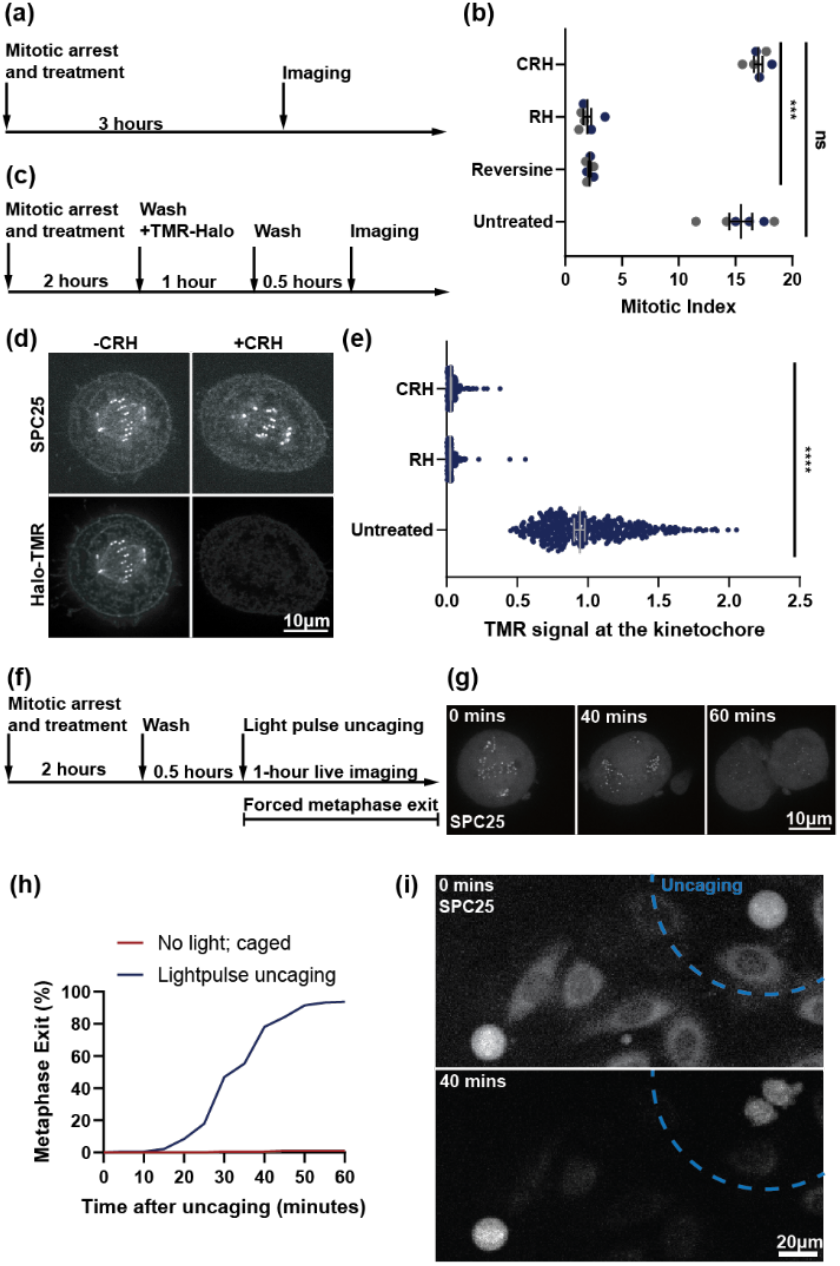
Light inducible spindle checkpoint silencing. (a-b) Checkpoint silencing assay. Cells expressing the Halo and GFP-tagged kinetochore protein SPC25 were arrested in mitosis with the CENP-E inhibitor GSK923295 (100 nM) in the absence (untreated) or presence of reversine (2 µM), reversine-halo (RH, 10 µM), or photocaged reversine-halo (CRH, 10 µM) for three hours, and then fixed to measure the proportion of mitotic cells (mitotic index). Each data point represents the mitotic index from 1000 cells; N = 6 measurements from three independent areas from two independent biological replicates; error bars: median with 95% CI; ***P ≤ 0.001. (c-e) Dye blocking assay. Cells expressing Halo and GFP-tagged SPC25 were arrested in mitosis by CENP-E inhibition in the absence (untreated) or presence of RH or CRH for 2 hours. Unbound probes were washed out, and cells were incubated with fluorescently labeled Halo ligand (Halo-TMR) for 30 min and then fixed for imaging (c). Representative images (d) show cells with or without CRH. Graph (e) shows Halo-TMR fluorescence intensity. Each data point represents the intensity from one kinetochore normalized to the mean of the untreated group (N = 337 - 414 kinetochores from 20 cells from two independent biological replicates); error bars: median with 95% CI; ****P ≤ 0.0001. (f-i) Cells stably expressing Halo and GFP-tagged SPC25 were arrested in mitosis by CENP-E inhibition and treated with CRH for 2 hours. After washing out unbound probes for 30 minutes, cells were exposed to a light pulse at t=0 to uncage CRH and imaged live to measure the time of anaphase onset. Representative images (g) show mitotic progression after uncaging. Graph (h) shows the distribution of waiting times until anaphase onset, with or without uncaging (N = 179 or 194, respectively). (i) Example images show local uncaging (blue dashed line) and checkpoint silencing with spatial precision.

To test whether CRH and RH bind to kinetochores as per our design, we carried out a dye-blocking assay^15^. In this assay, fluorescent-labeled Halo ligands stain the HaloTags that are not tethered by CRH or RH. (**Figure 3c**) Thus, CRH or RH binding should prevent fluorescent signals at kinetochores. In control cells arrested in mitosis by CENP-E inhibition, bright fluorescent signals from the halo ligands colocalize with kinetochores, as expected. In contrast, either CRH or RH treatment reduced signals at kinetochores by more than 20-fold, indicating their successful permeation into the cell and high occupancy at kinetochores. (**Figures 3d, e**)

We further tested if the photocage can be uncaged by a light pulse to release the kinetochore-localized reversine for SAC silencing. A successful photocage should allow mitotic exit only after light exposure. We loaded CRH into the cells arrested by CENP-E inhibition, washed out unbound CRH, exposed cells under a pulse of 385 nm light, and then measured mitotic timing by live imaging (**Figure 3f**). Cells without a light pulse stay arrested throughout the hour of observation, while most cells exposed to light exit mitosis between 20 and 50 minutes. These findings indicate that the light pulse successfully uncages CRH at kinetochores. (**Figures 3g, h**) CRH-PEG2 failed to induce checkpoint silencing after light activation (**Figure S4**), indicating that the linker was too short to allow the reversine motif to effectively engage MPS1. This result supports the requirement for a sufficiently long spacer to enable productive MPS1 binding.

To demonstrate one of the benefits of conditional localization pharmacology, we repeated the uncaging assay with the internal control in the same field of view. We imaged two cells simultaneously, but only one cell was exposed to light. Contrary to conventional reversine treatment, which would affect all mitotic cells, our localized uncaging induces mitotic exit specifically for the targeted cell (**Figure 3i**). This result illustrates the spatiotemporal precision of CLP. In these experiments, cells remained viable after the addition of CRH, as evidenced by the cells’ consistent ability to enter (Figure 3b) and exit (Figure 3h) mitosis with or without CRH.

Using CRH, we studied the late mitosis function of a kinesin-8 family motor protein, KIF18A, which regulates microtubule dynamics in mitosis^16, 17, 18^. KIF18A function before anaphase has been well studied, and deletion of kinesin-8 motors in fungi causes spindle hyper-elongation in late mitosis^19^, but SAC activation following KIF18A inhibition in mammalian cells has hampered studies of late mitosis. We used CRH to silence the SAC in individual cells following KIF18A knockdown, allowing us to track spindle dynamics after anaphase onset by live imaging with high temporal resolution (Figure 4a, b). Although KIF18A inhibition increased spindle length at anaphase onset, as previously reported in metaphase^16^, the spindle elongated more slowly in anaphase in KIF18A knockdown cells compared to control cells (**Figure 4c, e**). During cytokinesis, however, spindle length plateaued in control cells but continued to elongate in KIF18A knockdown cells (**Figure 4c, f**). This spindle elongation occurred in parallel with rapid cell elongation in cytokinesis (**Figure 4d**). We also found that formation of the spindle midbody, a cytokinesis microtubule structure derived from the central spindle, was disrupted in KIF18A knockdown cells (**Figure 4b**), likely due to disruption of the central spindle in anaphase as previously reported^20^.

**Figure 4.**
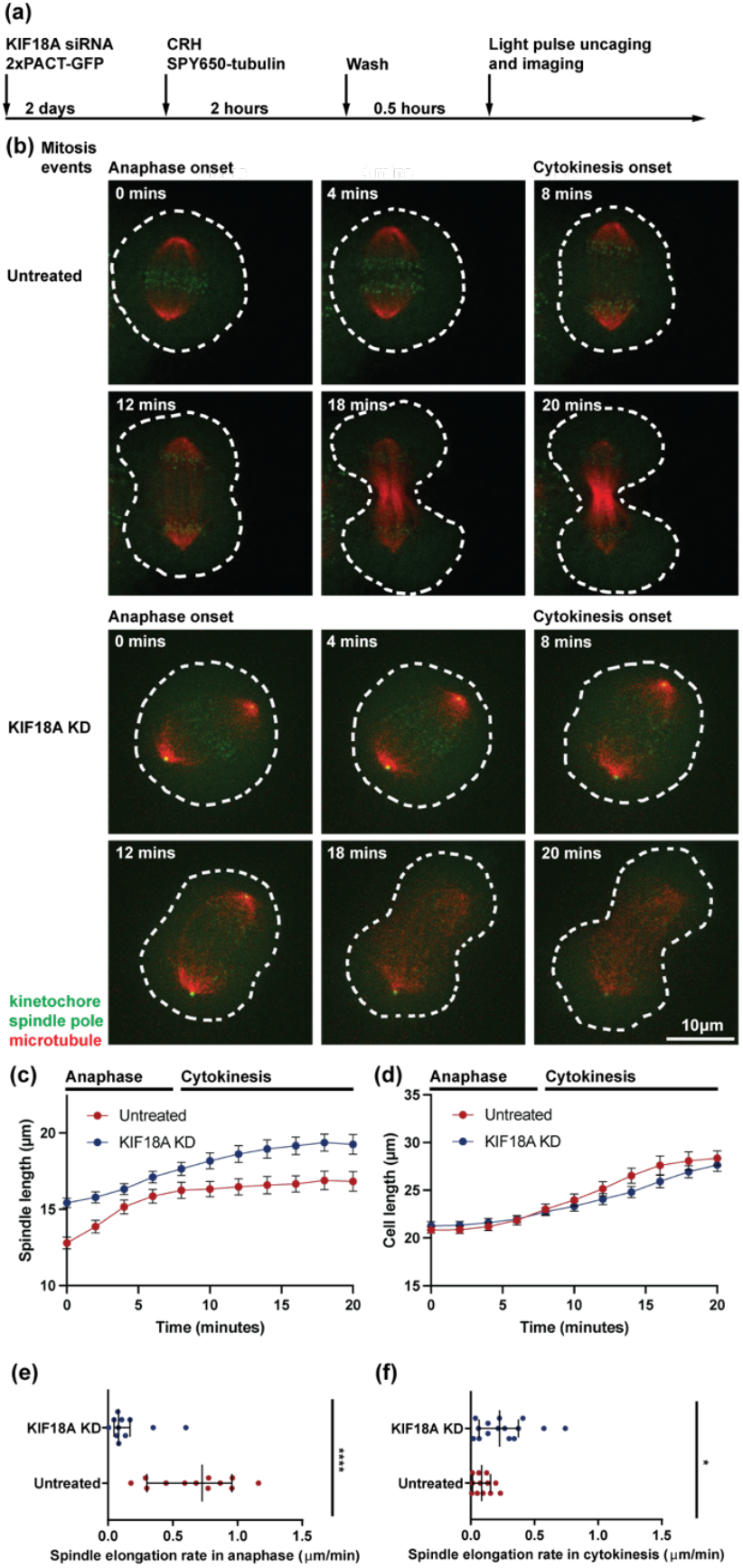
KIF18A regulates spindle elongation in late mitosis. Cells stably expressing the Halo- and GFP-tagged kinetochore protein SPC25 were treated with (KIF18A KD) or without (untreated) KIF18A siRNA (200 nM) for two days to knock down KIF18A. 2xPACT-GFP plasmid was transfected along with the siRNA to label the spindle poles. Cells were incubated with SPY650-tubulin (labels spindle microtubule) and CRH (10 μM) for two hours, before unbound probes were washed out. For each cell, CRH was uncaged with a light pulse, followed by live imaging. (a) Schematic of the experiment. (b) Example images show progression from anaphase onset (t = 0) to cytokinesis. (c-d) The lengths of the mitotic spindle (c) and the cell (d) during anaphase (t ~ 0-6 minutes) and cytokinesis (t ~ 8-20); N = 12 cells for untreated (red) or 15 cells for KIF18A KD (blue); error bars: mean and SEM. (e-f) Spindle elongation rates in anaphase (e) and cytokinesis (f); ****P ≤ 0.0001; * P ≤ 0.05; error bars: median and 95% CI.

Our results indicate that spindle elongation in late mitosis is restricted both by KIF18A and by the length of the cell. Spindles start longer in KIF18A knockdown cells at anaphase onset but elongate slowly relative to control cells until cell length increases to provide space. These dynamics suggest that the size of the cell limits the size of the spindle, rather than the spindle pushing to drive cell elongation^21^, with the caveat that disruption of the central spindle might compromise pushing forces exerted by the spindle in KIF18A knockdown cells. Finally, our findings suggest that KIF18A function in regulating spindle size and spindle midbody formation in late mitosis may contribute to the sensitivity of chromosomally unstable cancer cell lines to KIF18A inhibition^22^.

## CONCLUSIONS

We developed a SAC inhibition tool based on the CLP concept for conditional localized cell cycle control. Biological assays exhibited precise spatiotemporal control over spindle assembly checkpoint (SAC) silencing and mitotic exit. The use of CRH enabled us to achieve high time resolution, single-cell live imaging while silencing the Spindle Assembly Checkpoint to study the late mitosis function of KIF18A.

To extend the CLP design to broader application scenarios, proper understanding of how the ligand molecule is bound in the protein active site is critical. Chemical tethering (not limited to the Tag protein family) attachment points should be positioned toward clear exit points projecting from ligand binding sites to minimize the influence over ligand-protein binding interactions and may benefit greatly from already established structure activity relationships for certain protein classes. Photocages should be installed at key locations on ligands in a manner that will completely block critical interactions in the protein binding pocket. The photocage photophysical properties may also be tuned for uncaging by more red-shifted (lower energy) light or even using two-photon pathways. For ligand molecules bearing typical functional groups, such as hydroxy, amino, carboxylic acid, and thiol groups, photocaging is straightforward. However, in some cases, the optimal ligand screened for the target protein is typically “functional group saturated,” and no free or easily accessible functional group is available for photocaging. In these cases, less potent ligand molecules in the screening process could also provide a potential candidate for CLP modification because localization enhances the potency and working concentration by placing the target protein in proximity, providing the opportunity for enhanced potency and specificity. Biologically, the CLP concept could potentially benefit drug development for conventionally challenging targets. Creating anchoring ligands for endogenous proteins could pave the way for clinical application of CLP.

## ASSOCIATED CONTENT

### Supporting Information

A brief statement in nonsentence format listing the contents of material supplied as Supporting Information should be included, ending with “This material is available free of charge via the Internet at http://pubs.acs.org.”

## AUTHOR INFORMATION

### Notes

Any additional relevant notes should be placed here.

## ACKNOWLEDGMENT

We thank Dr. Jun Gu for NMR assistance, Dr. Charles W. Ross III for high-resolution mass spectrometry (HRMS) assistance. D.M.C. acknowledges NIH NIAMS (R01AR077094) and the University of Pennsylvania for funding. M.A. L. acknowledges NIH (GM122475 and P01-CA265794). G.Y.C. acknowledges Basser Center for BRCA grant.

