## Supporting Information for "Conditional Localization Pharmacology Manipulates Cell Cycle with Spatiotemporal Precision"

***General information***

All commercially available reagents and solvents were used as received.

6-chloro-1-bromohexane, NaH (60% in mineral oil), hexaethylene glycol and

N,N-diisopropylethylamine (DIEA) were purchased from TCI. *tert*-Butyl bromoacetate was purchased from Alfa Aesar. 6-Chloro-2-fluoro-9H-purine and 5-(1-Bromoethyl)-6-nitrobenzo[d][1,3]dioxole were purchased from Ambeed. Cyclohexylamine was purchased from thermal scientific. Cesium carbonate was purchased from Acros organic. PyBOP was purchased from Oakwood Chemical. All chemicals were used without further purification.

***Instrumentation***

- Thin-layer chromatography (TLC) was performed on Sorbent Technologies silica plates.
- Automated flash column chromatography was performed using RediSep Rf silica Gel or C18 column on CombiFlash Rf200 system with internal UV detector. The instrument is available from Teledyne Isco, Inc., NE., USA.
- Ultraviolet-Visible (UV-Vis) absorption spectrophotometry was performed on a JASCO V-650 spectrophotometer with a PAC-743R multichannel Peltier using quartz cells with a 1 cm cell path length.
- Proton nuclear magnetic resonance spectroscopy (1H NMR) and Carbon nuclear magnetic resonance spectroscopy (13C NMR) spectra were recorded on a Bruker NEO 600 MHz NMR and processed by MestReNova.
- Low-resolution mass spectra were obtained using Liquid-Chromatography-Mass-Spectrometry (LCMS) on Waters instrument, electrospray ionization in either positive or negative mode. High-resolution mass spectra (HRMS) were obtained at the University of Pennsylvania's Mass Spectrometry Service Center on Waters LC-TOF mass spectrometer (model LCT-XE Primer) using electrospray ionization in positive or negative mode, depending on the analytes. HRMS data analysis was performed using the automated Waters software.
- Preparatory highperformance liquid chromatography (HPLC) was performed on an Agilent 1290 Infinity II LC system equipped with a Phenomenex Luna Omega 5 μM PS C18 column using water and acetonitrile phase.

Note: Except for compound **1**. HRMS data for all the rest compounds is collected in Bruker scimaX instrument with mass accuracy (internal) up to 600 ppb. (https://www.bruker.com/en/products-and-solutions/mass-spectrometry/mrms/scimax.html)

***Cell Culturing***

HeLa RMCE acceptor cells[1] (obtained from E.V. Makayev, Nanyang Technological University, Singapore) stably expressing 3×Halo-GFP-SPC25[2, 3], were used for all cellular biology assays in the paper. Similar to previously described[2, 3], cells were cultured at 37°C, with humidified air added with 5% $CO_2$. Growth media consist of 10% FBS (Corning) and 1% penicillin/streptomycin (Life Technologies) in DMEM. 70% confluence is reached at the beginning of each assay.

***Plasmids***

For spindle pole labeling, the pcDNA3.1 backbone containing cytomegalovirus and T7 promoters was used to host two tandem copies of the PACT (Pericentrin and AKAP Centrosome Targeting)[4] domain fused to eGFP (2×PACT-GFP)[5].

***Microscopy***

For fixed-cell imaging, cells were plated on a poly-D-lysine (Sigma-Aldrich) coated, 22 x 22 mm glass coverslip (no. 1.5; Fisher Scientific) before treatments. At the end of the treatments, cells were washed with PBS before fixing. To fix the cells, the cells were incubated with 4% formaldehyde and 1% triton in PBS under 37°C for 20 minutes. Then, the cells were mounted to a glass microscope slide (Fisher Scientific) with VECTASHIELD® antifade mounting medium with DAPI (Vector Laboratories, Inc), and sealed with nail polish.

For live imaging, cells were incubated in L-15 medium without phenol red (Invitrogen) supplemented with 10% FBS (Corning) and 1% penicillin/streptomycin (Life Technologies) on FluoroDish™ (WPI). Throughout the experiment and imaging, the temperature was maintained at 37°C using an environmental chamber (Incubator BL; PeCon GmbH), and cells were imaged using a spinning disk microscope (DM4000; Leica) equipped with a 100× 1.4 NA oil immersion objective (Leica), an XY Piezo-Z stage (Applied Scientific Instrumentation), a CSU10 spinning disk (Yokogawa), an electron multiplier charge-coupled device camera (ImageEM; Hamamatsu Photonics), and a laser merge module (LMM5; Spectral Applied Research) equipped with 488- and 593-nm lasers, as previously described[2,3].

***Dye-blocking Assay***

Cells were arrested in metaphase using 100nM CENP-E inhibitor GSK923295 to facilitate sample collection with higher mitotic index. Meanwhile, the cells were also incubated with RH or CRH for 2 hours. At the end of the incubation, cells were then washed with PBS three times to remove excess RH or CRH. GSK923295 was added back with growth medium for continuing the metaphase arrest, and TMR-Halo (a fluorescent Halo dye) was added to stain for unoccupied Halo at the kinetochore over one hour. After the TMR-Halo staining, cells were washed with PBS, and fixed for imaging as described earlier in the Microscopy method.

***Mitotic Index Assay***

Cells were arrested with 100nM CENP-E inhibitor, GSK923295, and treated with reversine, RH, or CRH for three hours. The cells were then fixed, and mitotic index per 1000 cells were calculated under the microscope for two biological repeats each consisting of three mechanical repeats.

***Uncaging Assays***

Cells were arrested in metaphase using 100nM GSK923295 and treated with CRH for 2 hours to allow CRH binding to the kinetochore via 3Halo-GFP-SPC25. In some experiments, 10 µM verapamil was added to reduce efflux4 but effect was unperceivable. In uncaging assays performed under the same field of vision, cells were incubated for 3 hours instead of 2 to facilitate sample collection with higher mitotic index. After the incubation, the cells were washed three times with PBS to remove excess extracellular CRH, while GSK923295 was added back with growth medium to continue the metaphase arrest. The cells were rested for 30 minutes before uncaging to allow excess CRH to leave the cells. To uncage CRH, a 385nm light pulse was shone on all cells in the field of view or the indicated area within the field of view for 0.5 to 30 seconds. The difference in exposure was to accommodate for the optical differences in different magnifications and yielded similar results. The cells were then live imaged under the microscope for one hour. In this assay, metaphase was defined by the clear metaphase plate, depicted by the kinetochore SPC25-GFP signal as one column at the equator of the cell. Metaphase exist is defined by the initial separation of kinetochores on each sister chromatids.

***KIF18A Knockdown Assay***

200 nM siKIF18A (GCCAAUUCUUCGUAGUU-UU)[6] was transfected into the cells to knock down KIF18A, together with 300 ng 2×PACT-GFP plasmid to label the spindle poles, using Lipofectamine 3000® (Thermo Fisher Scientific).

Two days after transfection, cells were treated with 10 $\mu$M CRH for two hours

and then washed three times with PBS, as described above. Throughout the CRH treatment, SPY650-tubulin (Spirochrome) was added to the medium to label spindle microtubules. CRH was uncaged in one cell at a time using 385 nm light as above, and the cell was imaged every two minutes for the next 30-60 minutes.

***Data Analysis and Statistics***

In this paper, all images were processed with ImageJ, and z-stack signal max projection on selected stacks were adopted before further analysis. All the significance levels in the paper were obtained using Student T-test for two populations. For dye-blocking assay, TrackMate, an ImageJ plug-in, was used to define all particles with sizes close to a kinetochore. The quality, a parameter defined by the relative brightness and size in TrackMate, was used to gate the kinetochores from background particles in each image. The threshold is defined at the mean of quality + 3 standard deviations of quality in the image. The SPC25-GFP signal and the TMR signal at the kinetochore were calibrated against the mean of respective signal in the background for each picture. A ratio of calibrated TMR to SPC25-GFP signals was calculated per kinetochore. Then, the ratio from all kinetochores were pooled together for plotting. In the plot, the mean ratio of the control group was used to normalize all ratios for ease of understanding. For mitotic index assay, the mitotic index was calculated per 1000 cells, and mechanically repeated three times in different areas per one of

**Deleted:** In

**Commented [ML1]:** Unclear. When is the tubulin dye added? We want to give enough detailed information in this section for someone to be able to reproduce your protocol.

**Deleted:** meantime

**Deleted:** to

the two biological repeats. All six indexes per group were plotted as dots, while their mean and standard deviation being noted. For uncaging assays, experiments with various light exposure time and metaphase arrest time were pooled together to grant a large sample size. We spotted no sign of differences amongst all the groups pooled together.

For KIF18A knockdown assays, anaphase onset was defined by the initial separation of choromosomes, and cytokinesis onset was defined by the initial furrow ingression. Spindle length and cell length were measured along the mitotic axis of each cell every two minutes. To measure the rate of spindle elongation for each cell, spindle length was plotted against time, and rates were calculated as the slopes of two separate linear regressions on the anaphase and cytokinesis portions of the plot (Figure S5).

***In-vitro photolysis with Penn PhD Photoreactor M2***

CRH or CRH-PEG2 was dissolved in 1:1 MeCN/$H_2O$ to prepare a 1 mM solution (200 μL total). The solution was transferred to a 4 mL clear vial and irradiated in a Penn PhD Photoreactor M2 (365 nm, 20% intensity) with the fan set to 6800 rpm and stirring at 1000 rpm. Aliquots were withdrawn after 5, 10, and 20 min, and 10 μL of each was injected onto an analytical HPLC (LUNA OMEGA column) using a 35–50% MeCN/$H_2O$(0.1% TFA) gradient.

Commented [ML2]: correct?

Deleted: sister

Deleted: kineto

Deleted: e pairs

Deleted: .

Deleted: The average spindle length and cell length were plotted, with an error of SEM.

Deleted: The rates were plotted as dots, annotated with their median and 95% CI.

#### Supporting Figures

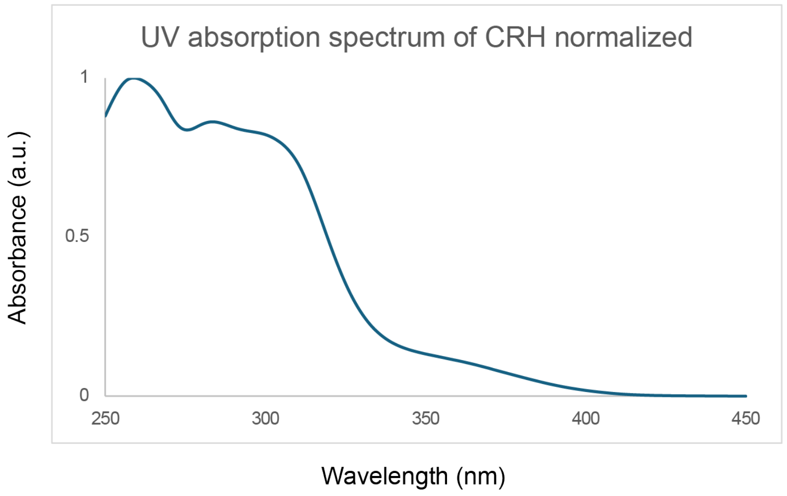


**Figure S1. Normalized UV absorption spectrum of CRH**

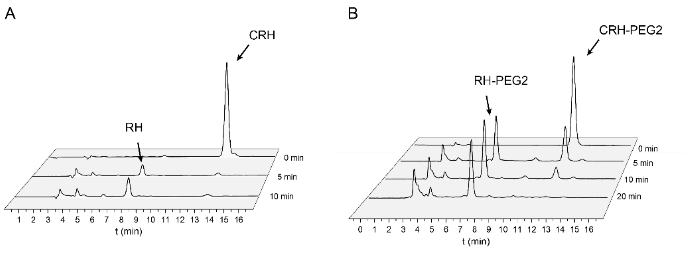


**Figure S2.** Analytical HPLC chromatograms of CRH uncaging in MeCN/$H_2O$. CRH was uncaged in PBS with 20% 365 nm light in Penn PhD Photoreactor M2. The products labelled in the chromatograms are recognized thanks to their mass and retention time. (A)CRH (B)CRH-PEG2

#### Synthesis and Characterization

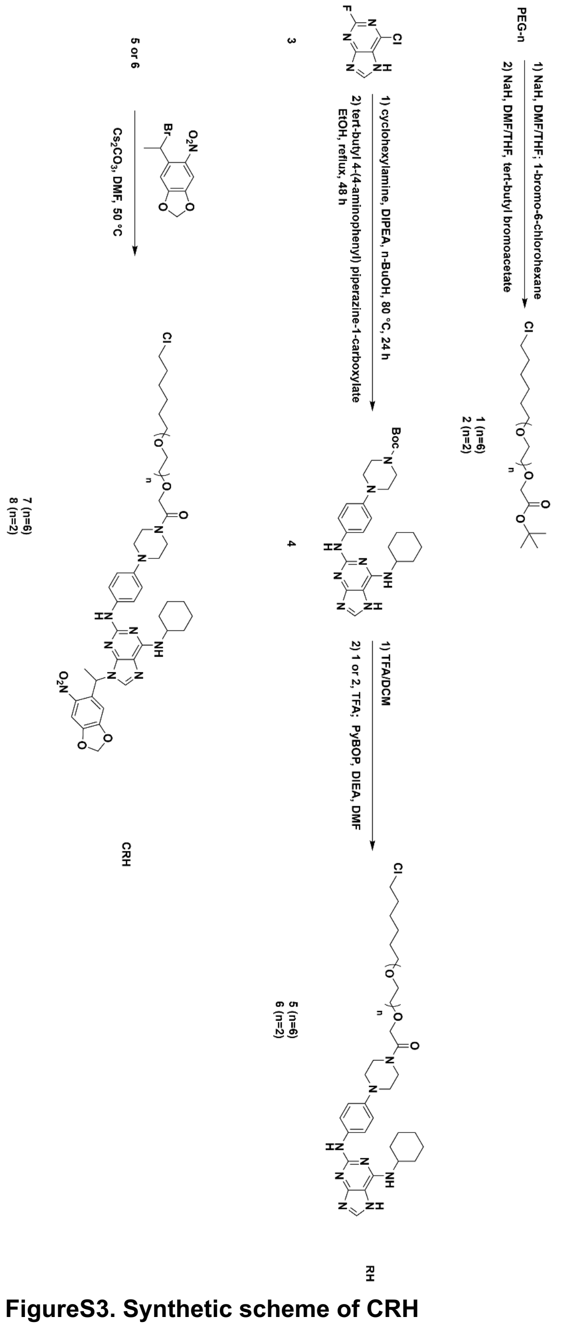


**FigureS3. Synthetic scheme of CRH**

*tert*-butyl 27-chloro-3,6,9,12,15,18,21-heptaoxaheptacosanoate(**1**)

Step 1: PEG-6(0.70 g, 2.47 mmol) was dissolved in 1:1 THF/DMF(2.5 M) and

sodium hydride (60% in mineral oil, 1.1 eq, 0.11 g) was slowly added into the mixture at 0 ºC. The reaction mixture was stirred at 0 ºC for 30 min and then 1-bromo-6-chloro hexane (1 eq, 0.37 mL) was added dropwise. The reaction was warmed to room temperature and stirred overnight. The reaction was quenched with sat. $NH_4Cl$ solution, diluted with 100 mL water and then extracted with ethylacetate(75 mL) twice. The organic phase was washed with brine, dried over sodium sulfate and then concentrated in vacuo. The mixture is directly used in the second step without further purification.
Step 2: Mixture(0.31 g) from step 1 was dissolved in 10 mL 1:1 THF/DMF and sodium hydride (60% in mineral oil, 1.1 eq, 34.3 mg) was slowly added into the mixture at 0 ºC. The reaction mixture was stirred at 0 ºC for 30 min and then *tert*-Butyl bromoacetate (1 eq, 140 μL) was added dropwise. The reaction was warmed to room temperature and stirred overnight. The reaction was quenched with sat. NH4Cl solution, diluted with 100 mL water and then extracted with ethylacetate (75 mL) twice. The organic phase was washed with brine, dried over sodium sulfate and then concentrated in vacuo. The crude mixture was subjected to column chromatography with a gradient of 0-3% methanol in DCM, yielding 0.15 g of 1 as a clear colorless oil (12% yield).

**$^1$H NMR** (600 MHz, $CDCl_3$) δ 4.01 (s, 2H), 3.73 – 3.66 (m, 4H), 3.66 – 3.61 (m, 18H), 3.57 (dd, *J* = 5.9, 3.9 Hz, 2H), 3.52 (t, *J* = 6.7 Hz, 2H), 3.45 (t, *J* = 6.6 Hz, 2H), 1.80 – 1.73 (m, 2H), 1.62 – 1.56 (m, 2H), 1.47 (s, 9H), 1.46 – 1.41 (m, 2H), 1.40 – 1.32 (m, 2H).

**$^{13}$C NMR** (151 MHz, $CDCl_3$) δ 169.80, 81.64, 71.36, 70.85, 70.75, 70.73, 70.72, 70.70, 70.24, 69.18, 45.17, 32.68, 29.59, 28.24, 26.83, 25.56.

**HRMS** (ESI, m/z): $C_{24}H_{47}ClO_9$ $[M+Na]^+$: 537.2831; found: 537.2806.

Following the same procedure, tert-butyl 2-(2-(2-((6-chlorohexyl)oxy)ethoxy) ethoxy)acetate **2** was obtained in 10% yield.

**$^1$H NMR** (600 MHz, $CDCl_3$) δ 4.02 (s, 2H), 3.74 – 3.67 (m, 4H), 3.65 (dd, *J* = 5.9, 3.9 Hz, 2H), 3.59 (dd, *J* = 5.9, 3.8 Hz, 2H), 3.53 (t, *J* = 6.7 Hz, 2H), 3.46 (t, *J* = 6.6 Hz, 2H), 1.77 (p, *J* = 6.9 Hz, 2H), 1.59 (dt, *J* = 14.8, 6.6 Hz, 3H), 1.47 (s, 9H), 1.46 – 1.42 (m, 2H), 1.37 (q, *J* = 7.2 Hz, 2H).
**$^{13}$C NMR** (151 MHz, $CDCl_3$) δ 169.84, 81.68, 71.39, 70.89, 70.84, 70.81, 70.78, 70.24, 69.21, 45.21, 32.70, 29.61, 28.26, 26.85, 25.58.
**HRMS** (ESI, m/z): $C_{16}H_{31}ClO_5$ $[M+Na]^+$: 361.175222; found: 361.175169.

tert-butyl4-(4-((6-(cyclohexylamino)-7H-purin-2-yl)amino)phenyl)-1$\lambda^4$-piperazine-1-carboxylate(**4**) was synthesized according to the reported procedure.[4]

Boc-reversine **4** (185 mg, 0.38 mmol) was dissolved in 2 mL 1:1 TFA/DCM and stirred for 30 min. Then the reaction mixture was concentrated in vacuo. PEG-6 halo **1** (193 mg, 0.38 mmol) dissolved in 2 mL 1:1 TFA/DCM and stirred for 30 min. Then the reaction mixture was concentrated in vacuo. Two mixture was combined and 20 eq DIEA was added to neutralize TFA residue. Then the mixture was diluted with 2 mL DMF and added PyBOP (195 mg, 0.38 mmol, 1eq). The reaction was stirred at room temperature for 30 min and then quenched with sat. $NH_4Cl$ solution. The resultant solution was diluted with 50 mL water and then extracted with ethylacetate(30 mL) twice. The organic phase was washed with brine, dried over sodium sulfate and then concentrated in vacuo. The crude mixture was subjected to column chromatography with a gradient of 5-10% methanol in DCM, yielding 177 mg of **5** as a brown oil (56% yield).

**$^1$H NMR** (600 MHz, MeOD) δ 7.74 (s, 1H), 7.60 (s, 2H), 6.93 (s, 2H), 4.32 – 4.25 (m, 2H), 4.10 (s, 1H), 3.73 (t, *J* = 5.0 Hz, 2H), 3.70 – 3.55 (m, 24H), 3.52 (ddd, *J* = 8.8, 6.1, 2.6 Hz, 4H), 3.45 – 3.39 (m, 2H), 3.09 (s, 2H), 2.13 – 2.06 (m, 2H), 1.83 (dt, *J* = 13.5, 3.6 Hz, 2H), 1.75 – 1.65 (m, 3H), 1.57 – 1.50 (m, 2H), 1.49 – 1.26 (m, 10H).
**$^{13}$C NMR** (151 MHz, MeOD) δ 170.10, 158.65, 154.98, 152.50, 147.25, 137.54, 136.39, 121.57, 118.81, 113.87, 72.14, 71.54, 71.51, 71.48, 71.45, 71.42, 71.40, 71.38, 71.35, 71.32, 71.30, 71.12, 71.09, 70.80, 70.78, 54.82, 52.13, 51.61, 50.46, 49.57, 46.10, 45.73, 43.04, 34.08, 33.75, 30.54, 27.73, 26.86, 26.49, 26.19.
**HRMS** (ESI, m/z): $C_{41}H_{65}ClN_8O_8$ $[M+H]^+$: 833.471296; found: 833.468665.

Following the same procedure, RH-PEG2 **6** was obtained in 70% yield.

**$^1$H NMR** (600 MHz, MeOD) δ 8.10 (s, 1H), 7.48 (s, 2H), 7.06 (s, 2H), 4.31 (s, 2H), 4.05 (s, 1H), 3.78 – 3.75 (m, 2H), 3.74 – 3.68 (m, 4H), 3.68 – 3.66 (m, 2H), 3.62 (dd, *J* = 6.1, 3.3 Hz, 2H), 3.56 (dd, *J* = 5.6, 3.6 Hz, 2H), 3.51 (t, *J* = 6.6 Hz, 2H), 3.44 (td, *J* = 6.6, 1.1 Hz, 2H), 3.23 (d, *J* = 21.5 Hz, 3H), 2.12 – 2.05 (m, 2H), 1.84 (q, *J* = 4.2 Hz, 2H), 1.75 – 1.67 (m, 3H), 1.59 – 1.52 (m, 2H), 1.48 – 1.23 (m, 10H).
**$^{13}$C NMR** (151 MHz, MeOD) δ 170.20, 153.02, 149.58, 148.44, 142.05, 131.80, 124.90, 118.41, 107.95, 72.19, 71.68, 71.53, 71.44, 71.15, 70.97, 52.13, 51.33, 50.85, 45.94, 45.71, 42.81, 33.74, 33.37, 30.55, 30.54, 27.74, 26.53, 26.49, 26.12.

**HRMS** (ESI, m/z): $C_{33}H_{49}ClN_8O_4$ $[M+H]^+$:657.363806; found: 657.364345.
Reversine Halo **5**(138 mg, 0.16 mmol), 5-(1-Bromoethyl)-6-nitrobenzo[d][1,3] Dioxole (90 mg, 0.32 mmol) and cesium carbonate(108 mg, 0.32 mmol) were

added in 2 mL DMF. The reaction was stirred overnight at 60 ºC and then quenched with sat. NH4Cl solution. The resultant solution was diluted with 30 mL water and then extracted with ethylacetate(20 mL) twice. The organic phase was washed with brine, dried over sodium sulfate and then concentrated in vacuo. The crude mixture was purified by prep HPLC with ingredient 40-55% acetonitrile:water, yielding a yellow oil product **7**(25 mg, 15% yield). The product peak comes up around 53% acetonitrile:water.

**$^{1}$H NMR** (600 MHz, $CDCl_3$) δ 9.06 – 9.04 (m, 1H), 8.50 (d, J = 9.7 Hz, 1H), 7.95 (s, 1H), 7.55 (s, 1H), 7.27 (d, J = 6.9 Hz, 2H), 6.99 – 6.94 (m, 2H), 6.49 (s, 1H), 6.30 (q, J = 6.9 Hz, 1H), 6.08 (d, J = 29.9 Hz, 2H), 4.70 – 4.63 (m, 1H), 4.30 (s, 2H), 3.83 (s, 2H), 3.76 – 3.70 (m, 2H), 3.70 – 3.61 (m, 22H), 3.56 (dd, J = 5.9, 3.6 Hz, 2H), 3.52 (t, J = 6.7 Hz, 2H), 3.44 (t, J = 6.7 Hz, 2H), 3.23 (d, J = 15.7 Hz, 4H), 2.13 – 2.05 (m, 2H), 2.02 (d, J = 7.0 Hz, 3H), 1.88 – 1.80 (m, 2H), 1.79 – 1.72 (m, 2H), 1.67 (d, J = 13.1 Hz, 1H), 1.62 – 1.54 (m, 2H), 1.52 – 1.40 (m, 6H), 1.39 – 1.31 (m, 2H), 1.31 – 1.24 (m, 1H).
**$^{13}$C NMR** (151 MHz, $CDCl_3$) δ 168.22, 152.87, 152.65, 149.23, 148.76, 147.84, 146.64, 142.06, 138.02, 133.56, 131.41, 121.74, 117.88, 112.56, 106.02, 105.89, 103.56, 71.32, 70.52, 70.40, 70.33, 70.27, 70.23, 70.18, 70.07, 69.91, 53.30, 50.99, 50.66, 50.39, 45.20, 44.53, 41.63, 33.14, 33.11, 32.65, 29.53, 26.81, 25.52, 25.15, 24.62, 20.72.
**HRMS** (ESI, m/z): $C_{50}H_{72}ClN_9O_{12}$ [M+Na]$^+$: 1048.48964; found: 1048.48812.

Following the same procedure, CRH-PEG2 **8** was obtained in 22% yield.

**$^{1}$H NMR** (600 MHz, $CDCl_3$) δ 9.22 (s, 1H), 8.42 (d, *J* = 8.6 Hz, 1H), 7.96 (s, 1H), 7.56 (s, 1H), 7.32 (d, *J* = 8.5 Hz, 2H), 7.07 (d, *J* = 8.5 Hz, 2H), 6.49 (s, 1H), 6.30 (q, *J* = 6.9 Hz, 1H), 6.12 – 6.05 (m, 2H), 4.67 (s, 1H), 4.28 (s, 2H), 3.89 (s, 2H), 3.72 (s, 6H), 3.67 (d, *J* = 4.5 Hz, 2H), 3.60 (s, 2H), 3.55 – 3.45 (m, 6H), 3.29 (d, *J* = 11.1 Hz, 4H), 2.09 (s, 2H), 2.03 (d, *J* = 7.0 Hz, 3H), 1.83 (d, *J* = 8.6 Hz, 2H), 1.75 (dt, *J* = 14.7, 6.8 Hz, 2H), 1.67 (d, *J* = 11.9 Hz, 1H), 1.59 (p, *J* = 7.1 Hz, 2H), 1.52 – 1.39 (m, 6H), 1.34 (q, *J* = 7.9 Hz, 2H), 1.30 – 1.24 (m, 1H).
**$^{13}$C NMR** (151 MHz, $CDCl_3$) δ 168.22, 152.87, 152.56, 149.15, 148.76, 147.89, 144.92, 142.12, 138.12, 133.42, 132.87, 121.76, 118.67, 112.65, 105.98, 105.93, 103.57, 71.57, 70.46, 70.34, 70.03, 69.75, 69.67, 53.36, 51.44, 51.15, 51.02, 45.19, 44.15, 41.30, 33.15, 33.12, 32.63, 31.08, 29.25, 26.76, 25.38, 25.15, 24.62, 20.75.
**HRMS** (ESI, m/z): $C_{42}H_{56}ClN_9O_8$ [M+Na]$^+$: 850.401314; found: 850.400550.

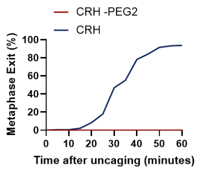


**Figure S4. PEG linker length affects reversine activity after uncaging.** Cells were treated with 10 µM CRH or its 2-PEG linker version (CRH-PEG2) following the experimental procedure identical to Figure 3f. Uncaging of CRH-PEG2 did not cause cells to exit metaphase, while uncaging CRH caused cells to exit metaphase within an hour.

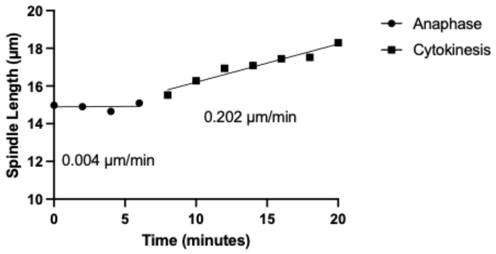


**Figure S5. Example analysis of spindle elongation rate.** Spindle length vs time is plotted for a KIF18A knockdown cell, with regression lines plotted for the anaphase and cytokinesis portions. The slopes are the spindle elongation rates.

Commented [ML5]: This figure would be nicer as one panel with both regression lines shown in the first panel, rather than having two additional plots. Also, I think you can just give the slope in um/min rather than writing out the equation for the line.

### NMR spectra

Deleted: ¶

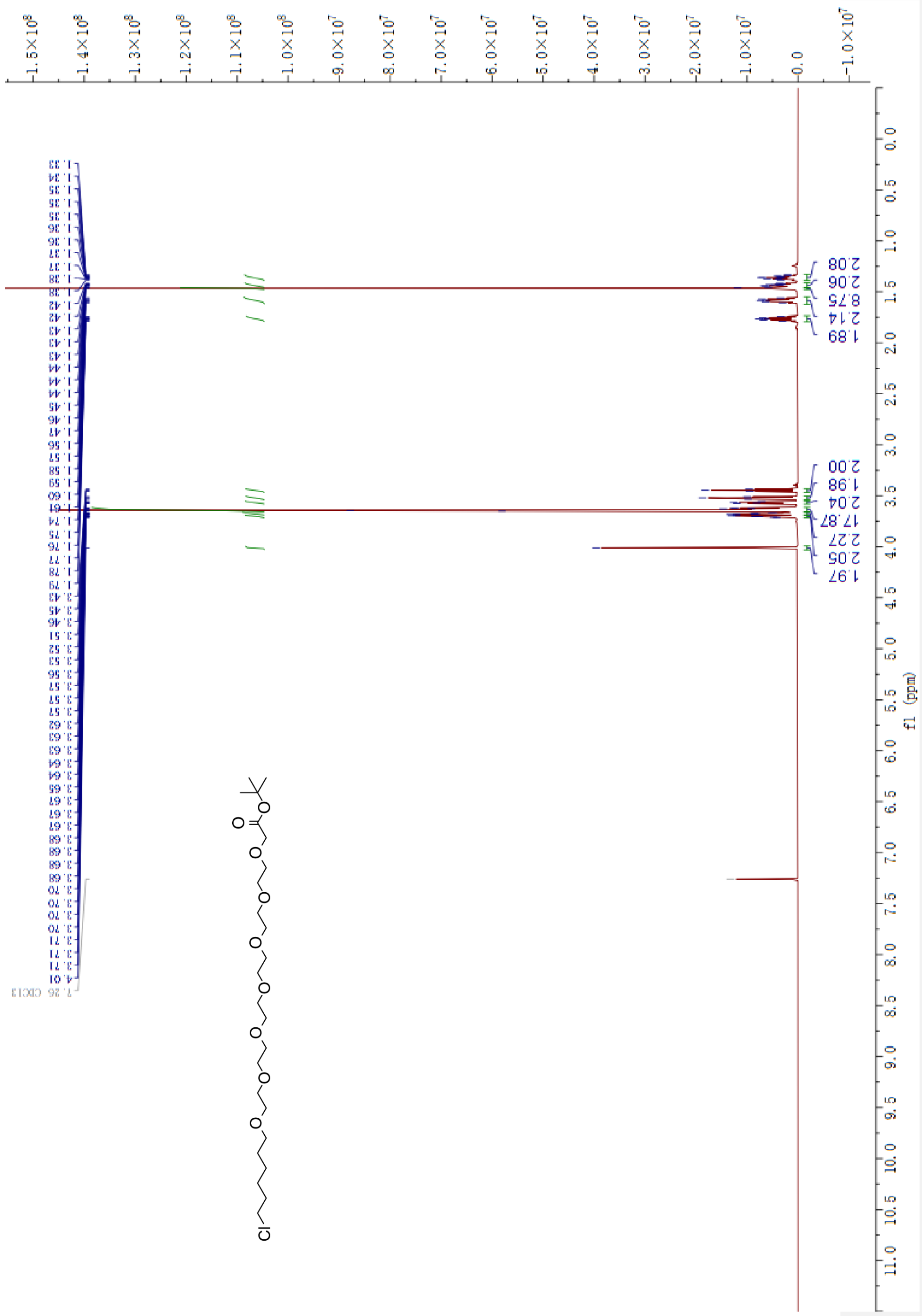


$^{1}H$ NMR spectrum of **1**(PEG-6 halo) in $CDCl_3$(600 MHz)

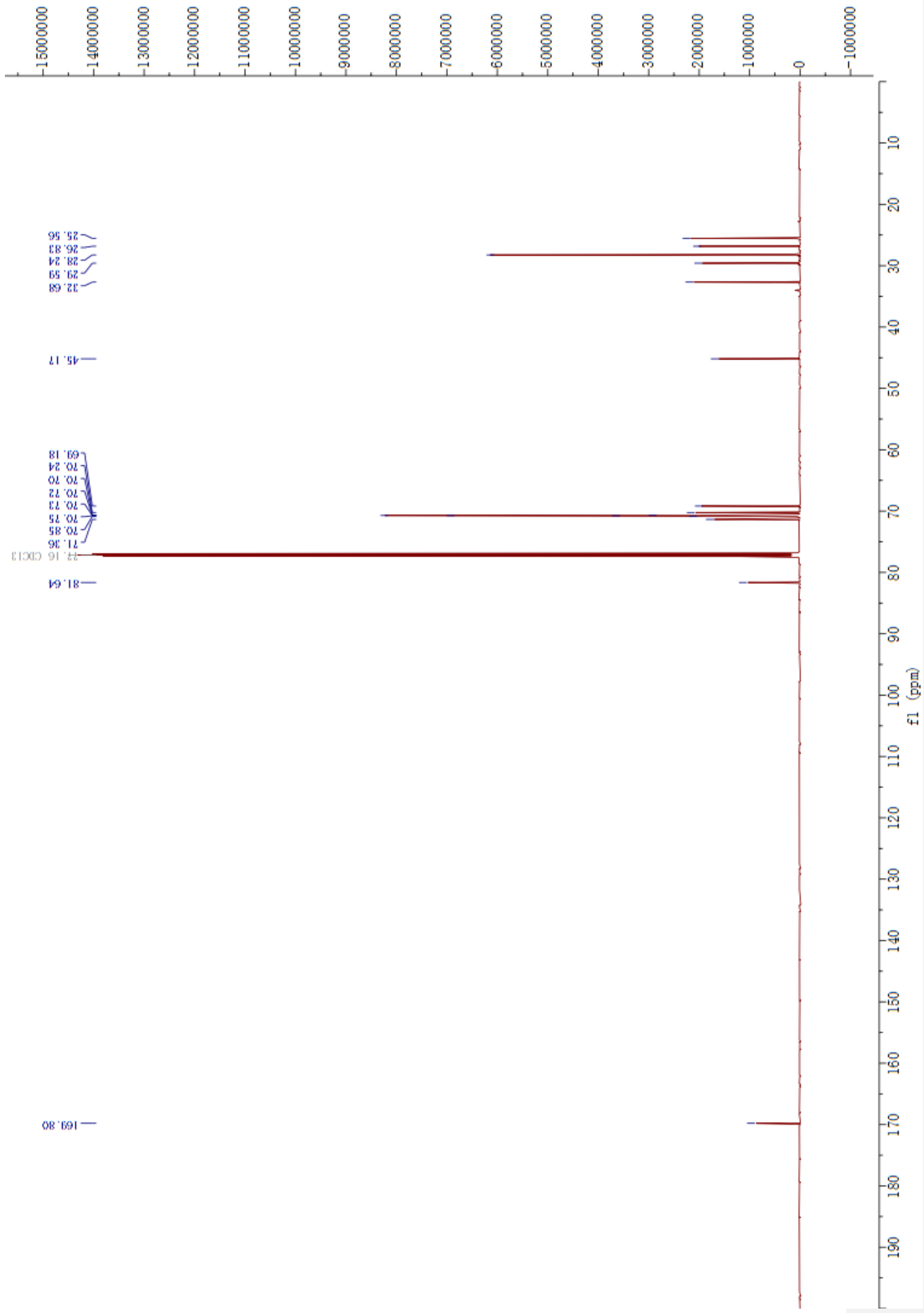


$^{13}$C NMR spectrum of **1**(PEG-6 halo) in $CDCl_3$(151 MHz)

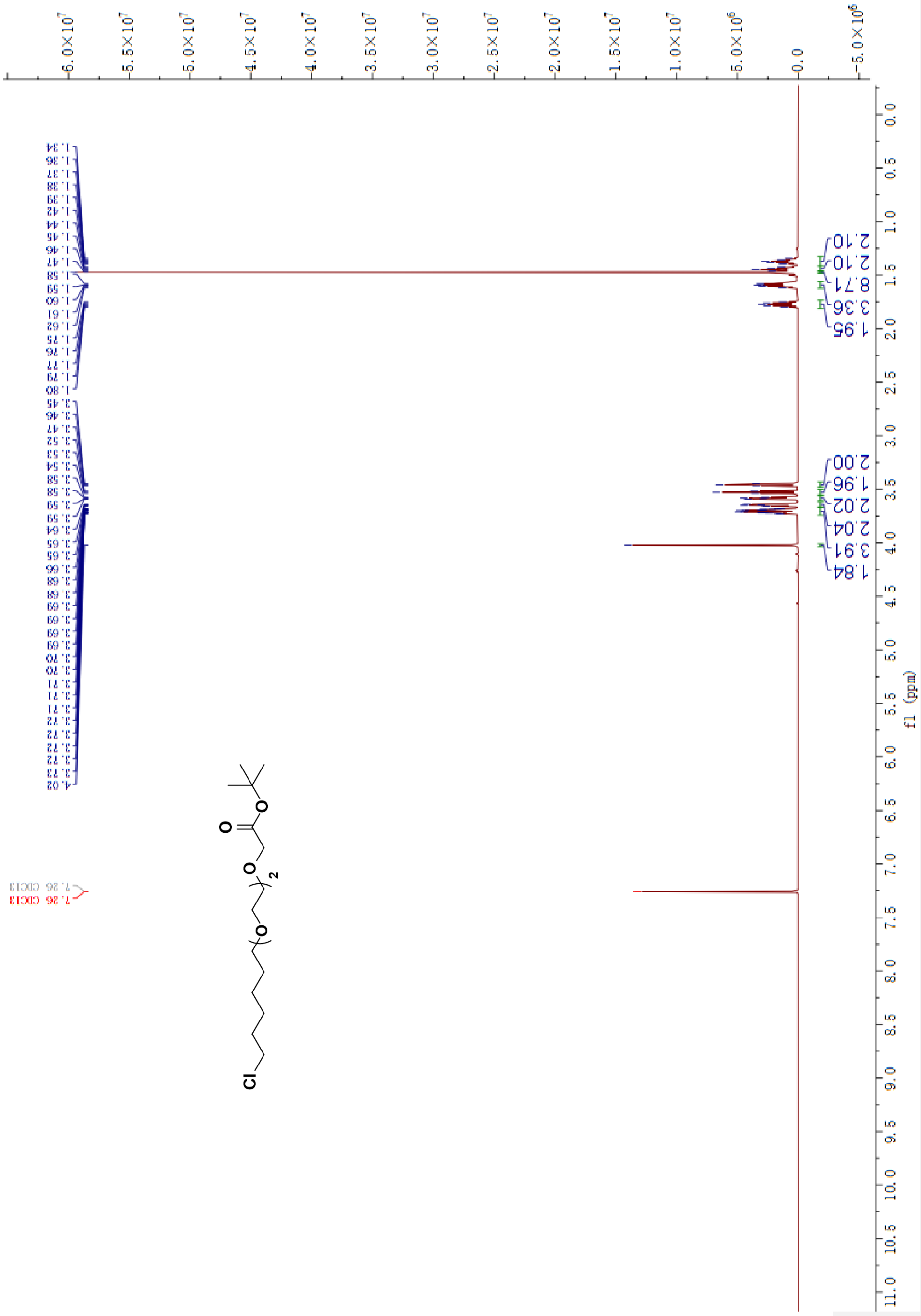


$^{1}H$ NMR spectrum of **2**(PEG-2 halo) in $CDCl_3$(600 MHz)

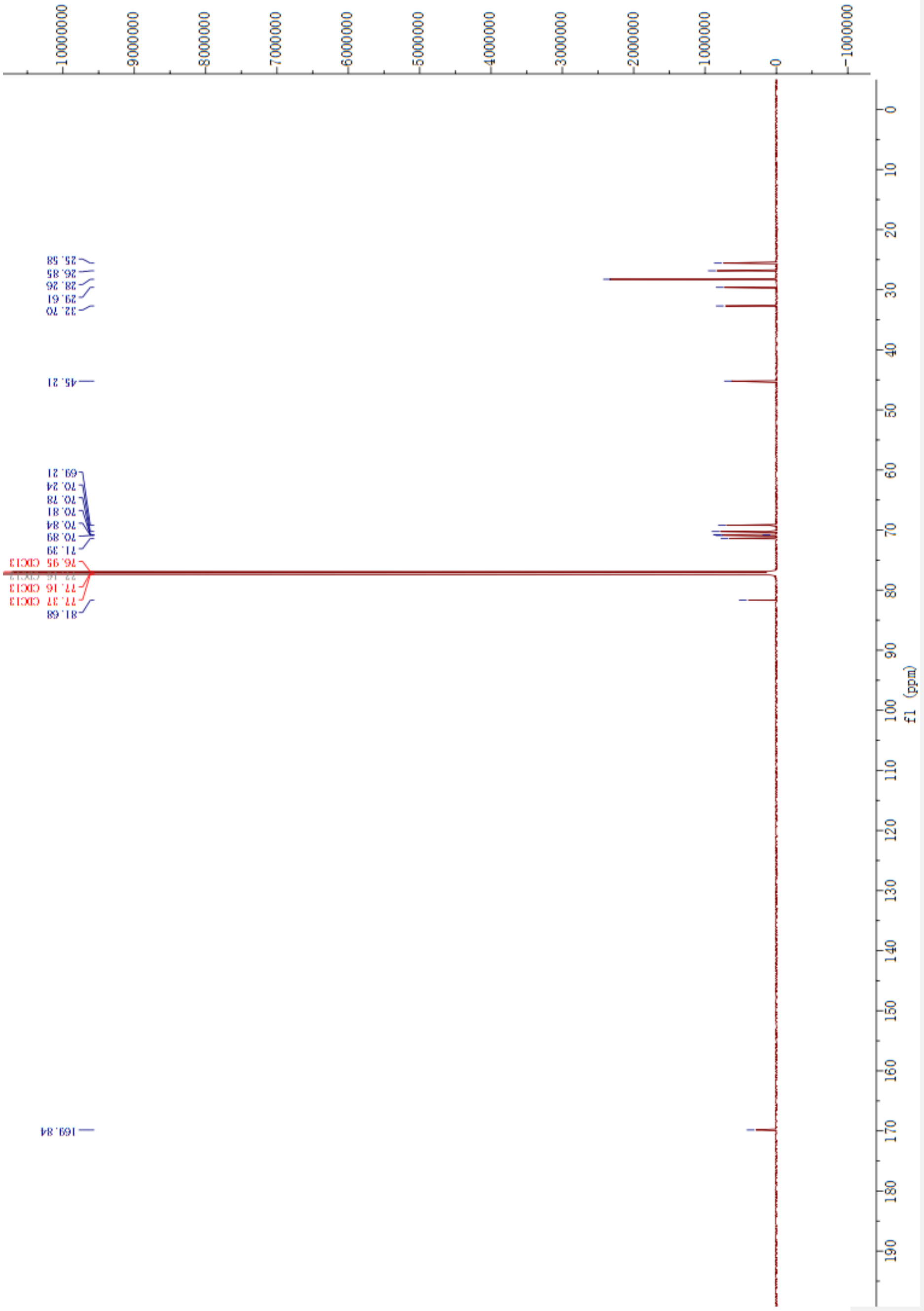


$^{13}C$ NMR spectrum of **2**(PEG-2 halo) in $CDCl_3$(151 MHz)

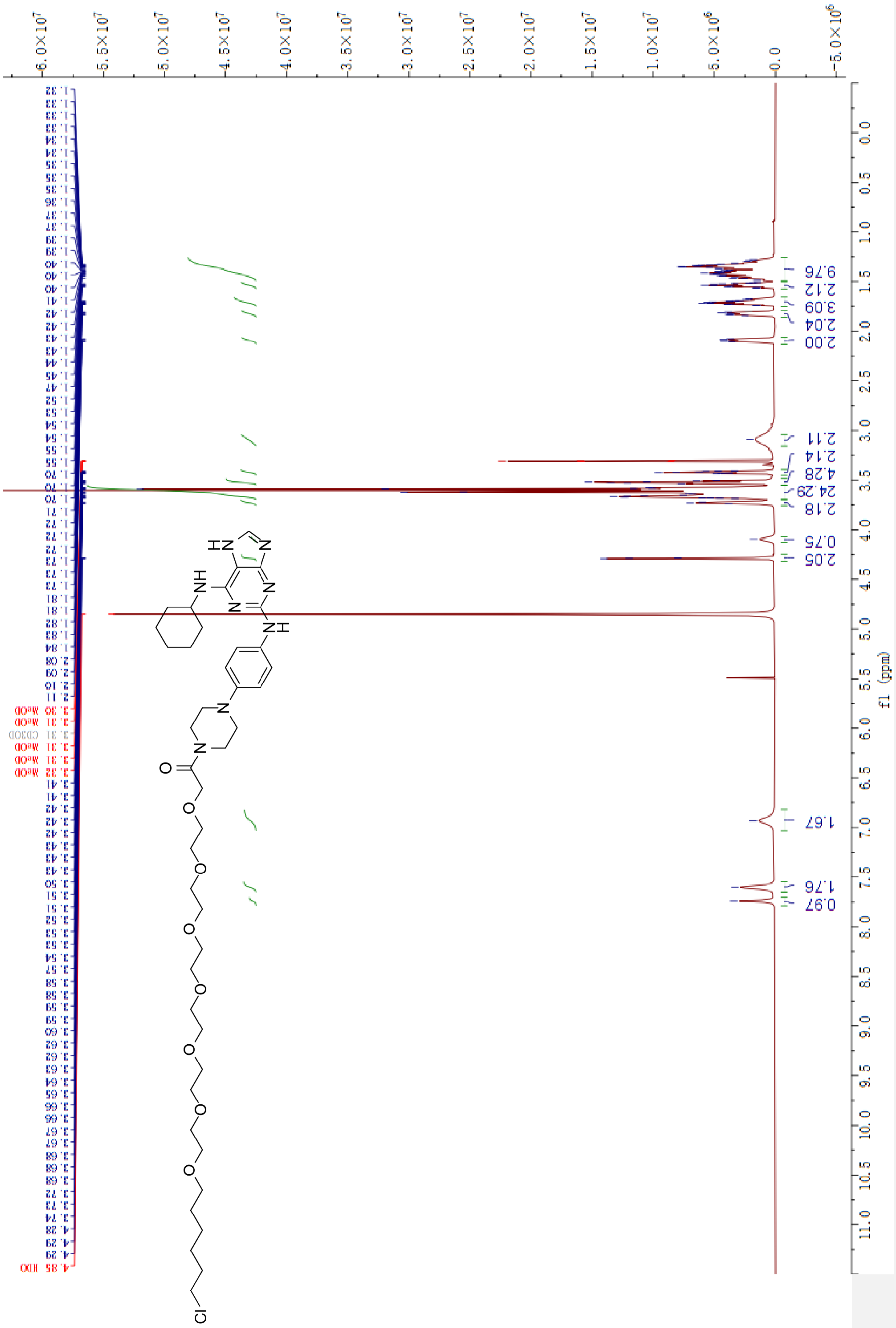

[1]H NMR spectrum of **5**(RH) in MeOD-d[4](600 MHz)

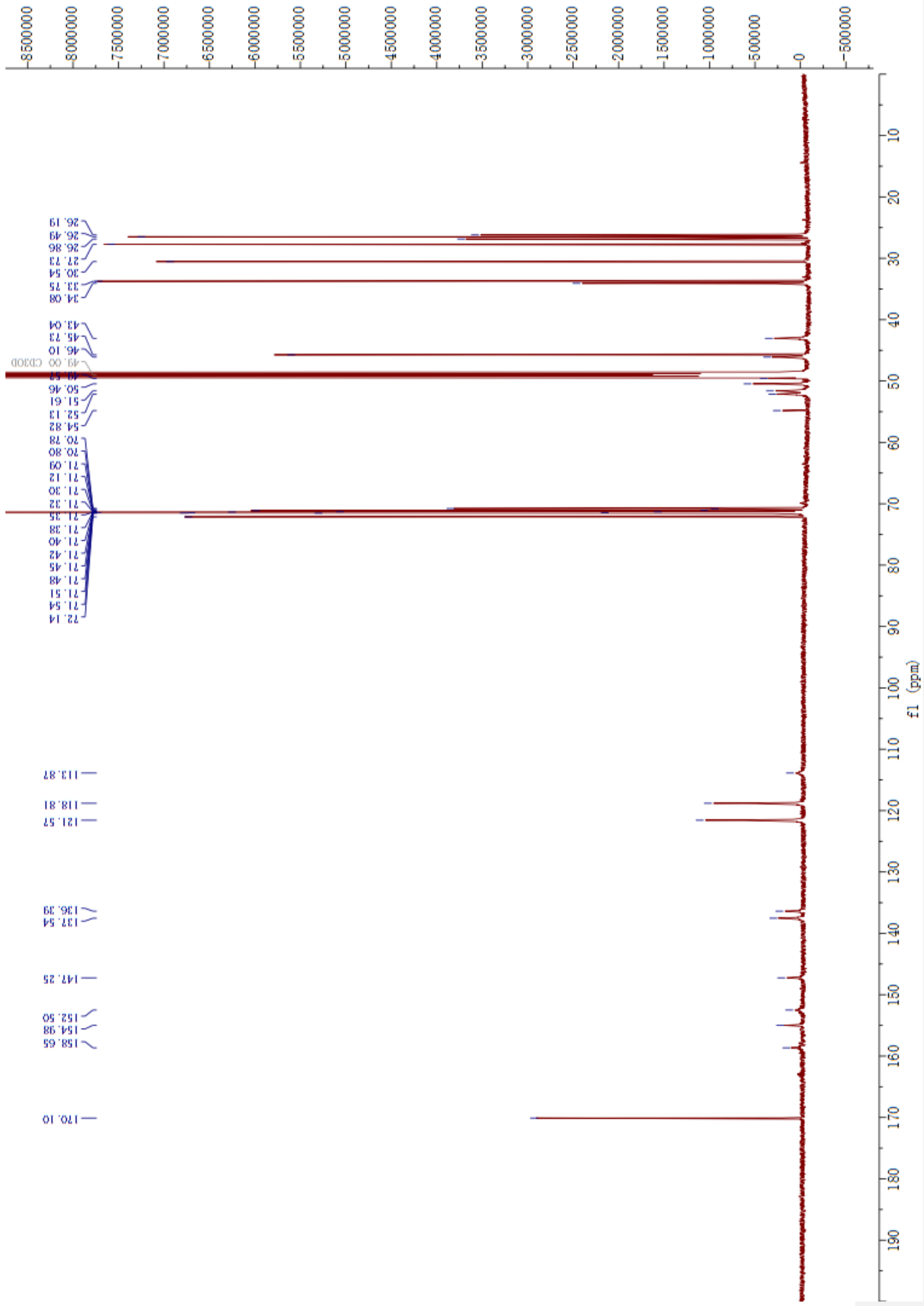


$^{13}$C NMR spectrum of **5**(RH) in MeOD-d$^4$(151 MHz)

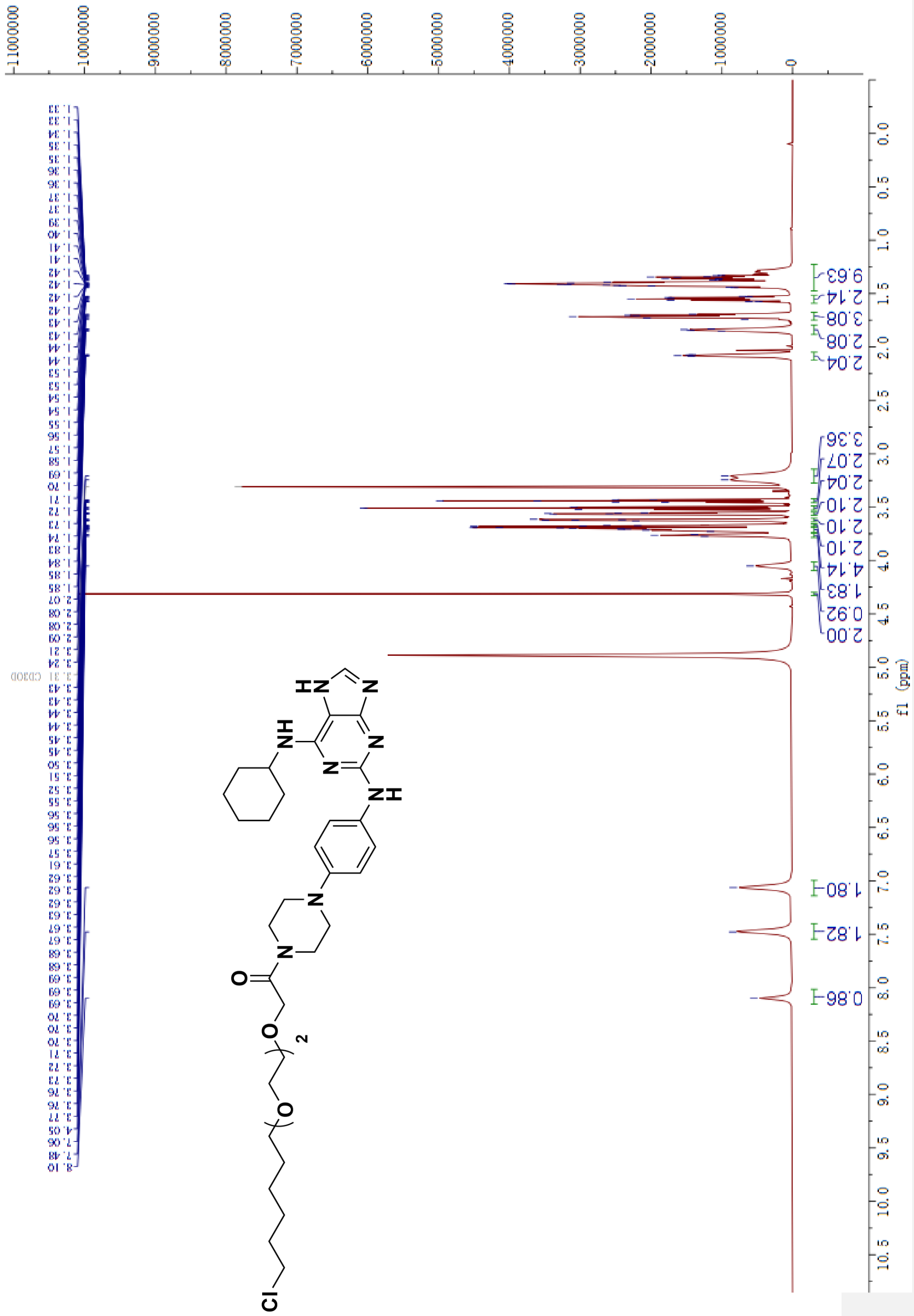

$^{1}$H NMR spectrum of **6**(RH-PEG2) in MeOD-d$^{4}$(600 MHz)

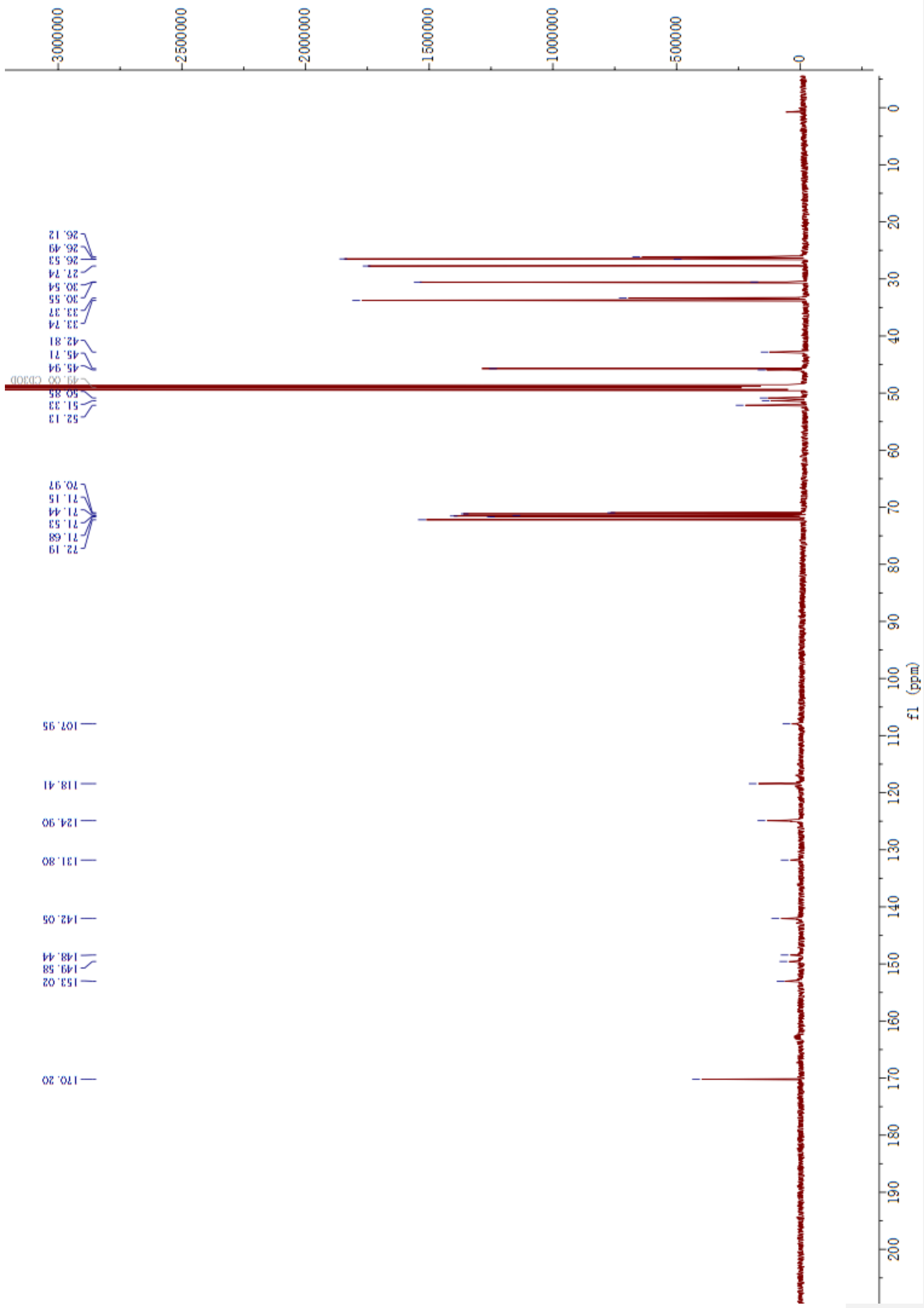


$^{13}C$ NMR spectrum of **6**(RH-PEG2) in MeOD-$d^4$(151 MHz)

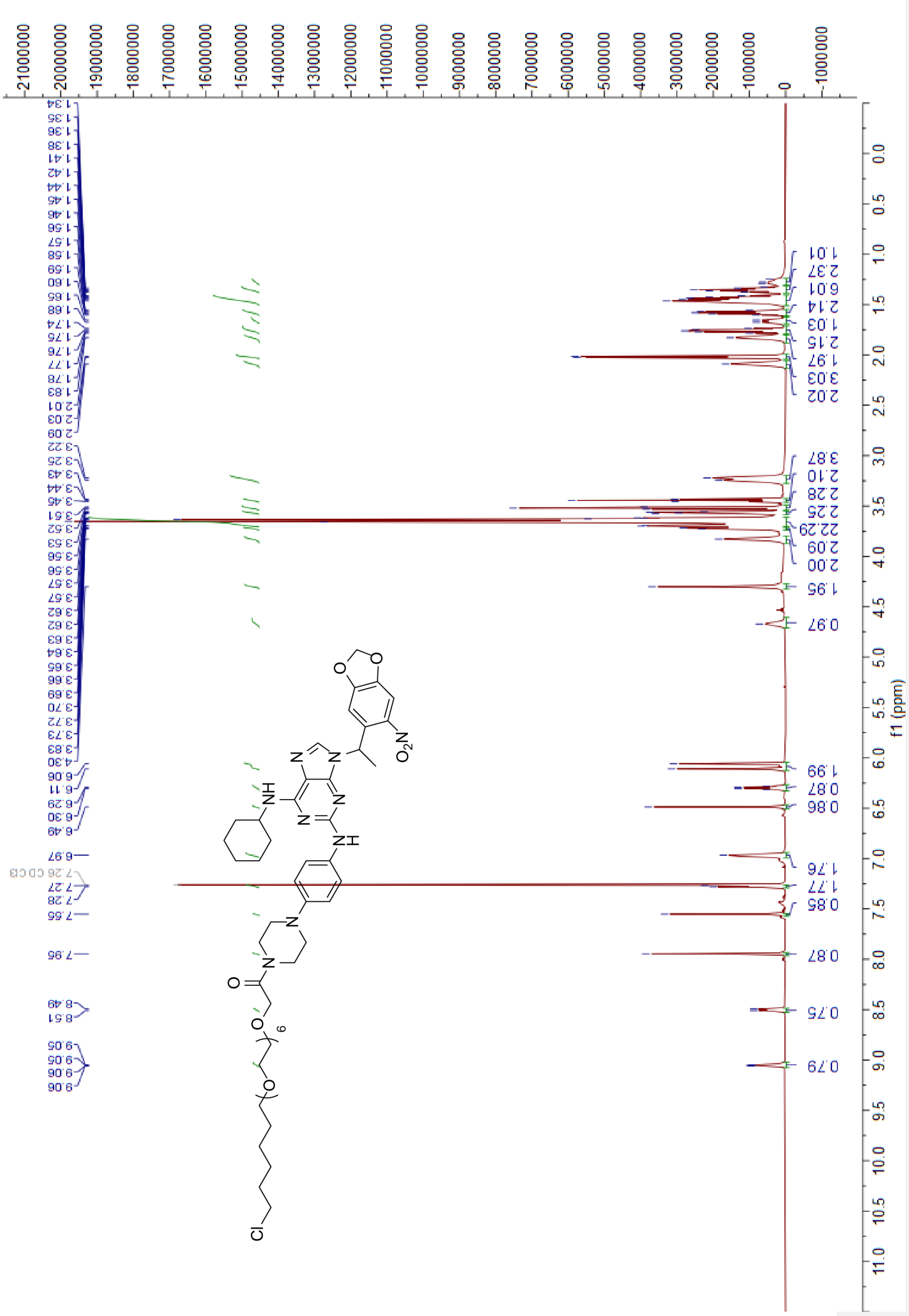

$^{1}H$ NMR spectrum of **7**(CRH) in $CDCl_3$(600 MHz)

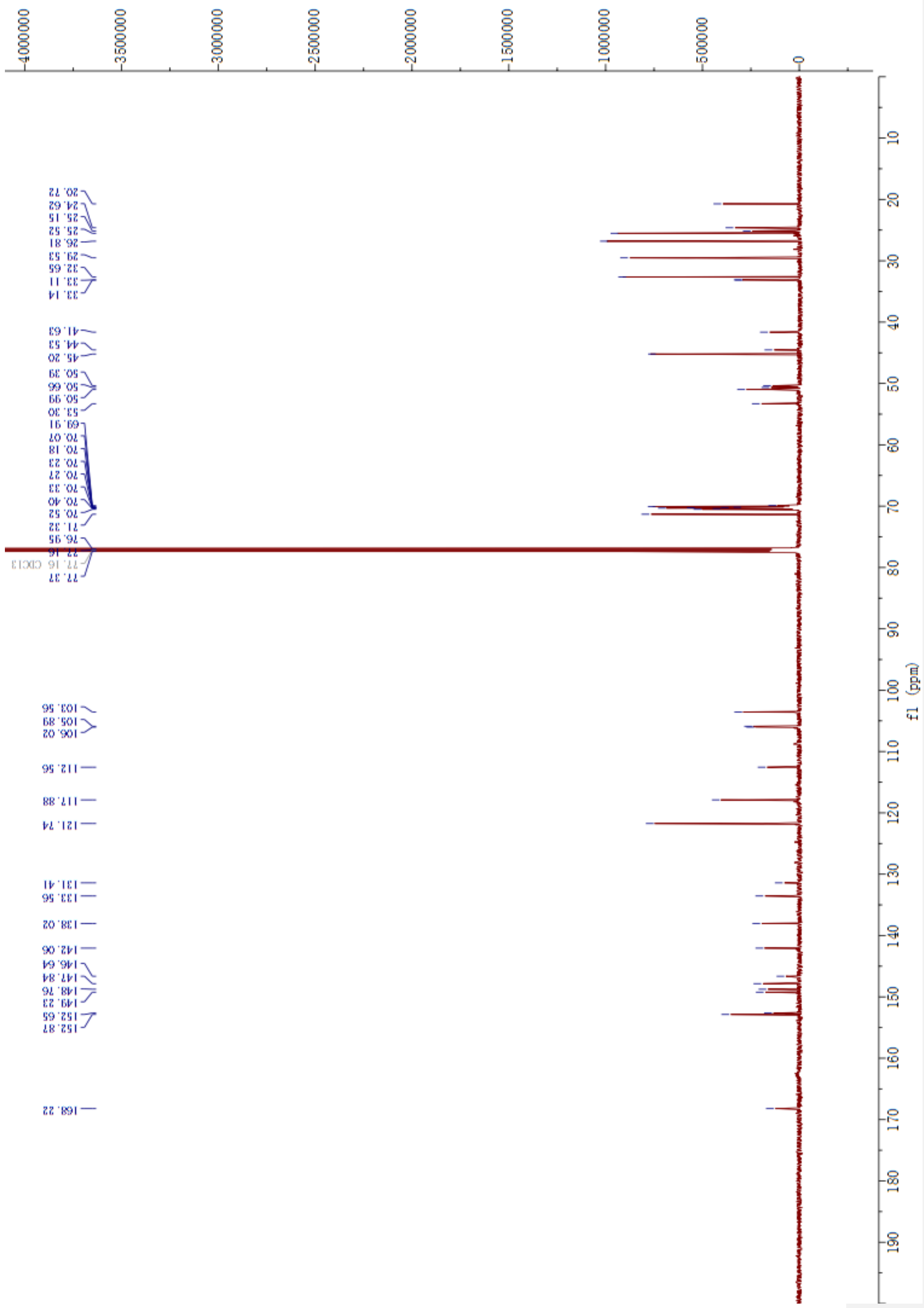


$^{13}C$ NMR spectrum of **7**(CRH) in $CDCl_3$(151 MHz)

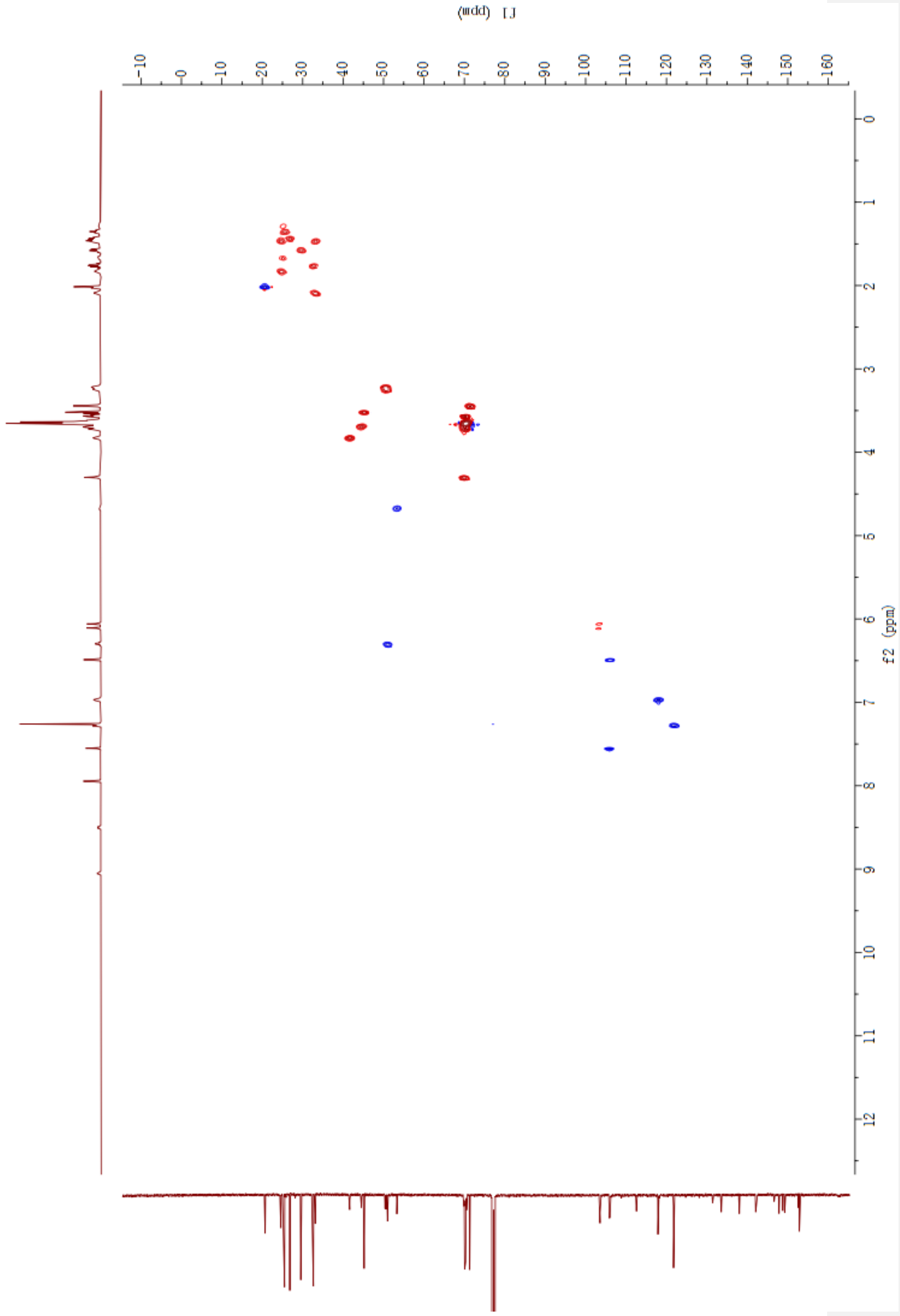


HSQC spectrum of **7**(CRH) in $CDCl_3$

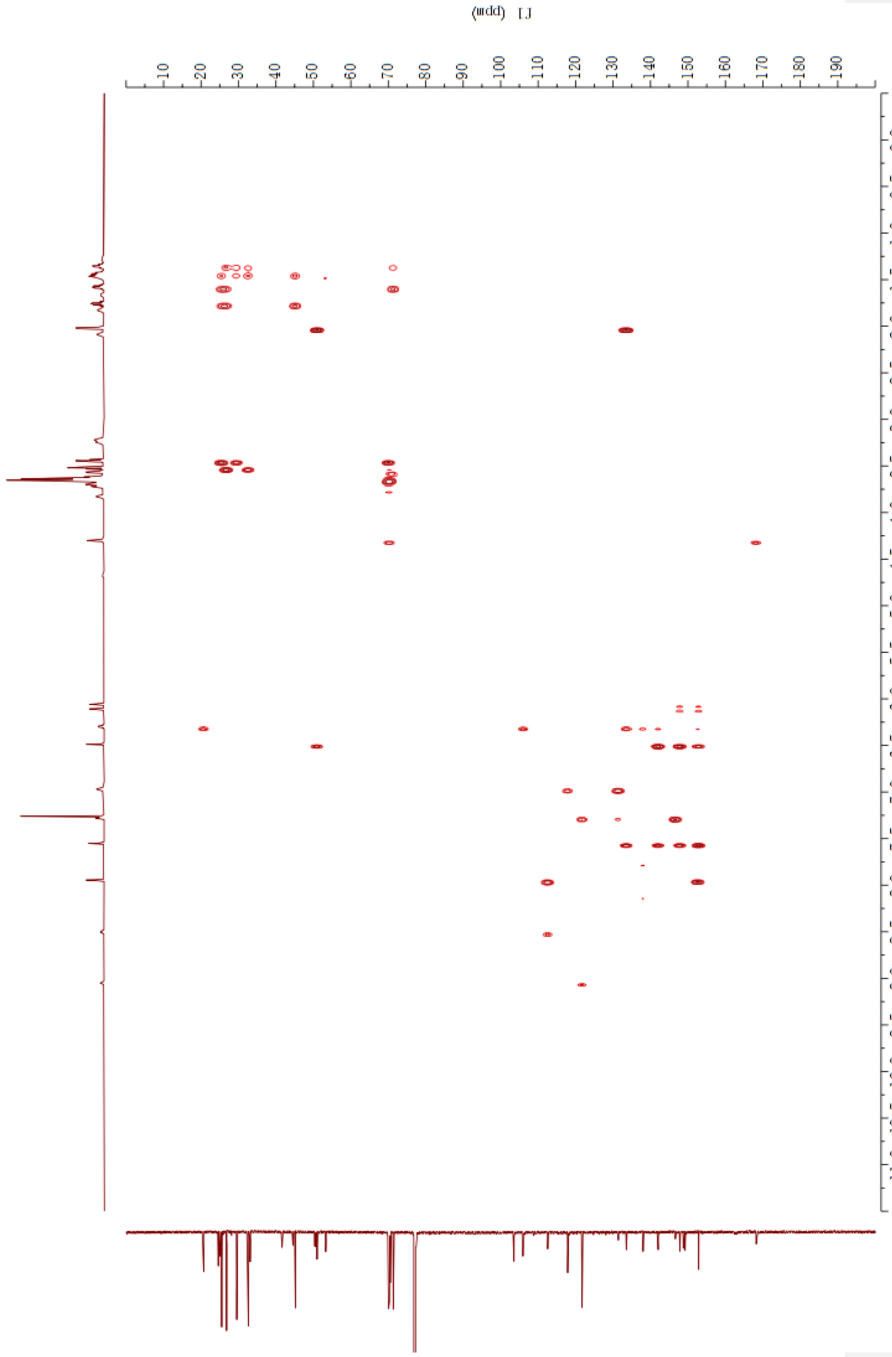

HMBC spectrum of **7**(CRH) in $CDCl_3$

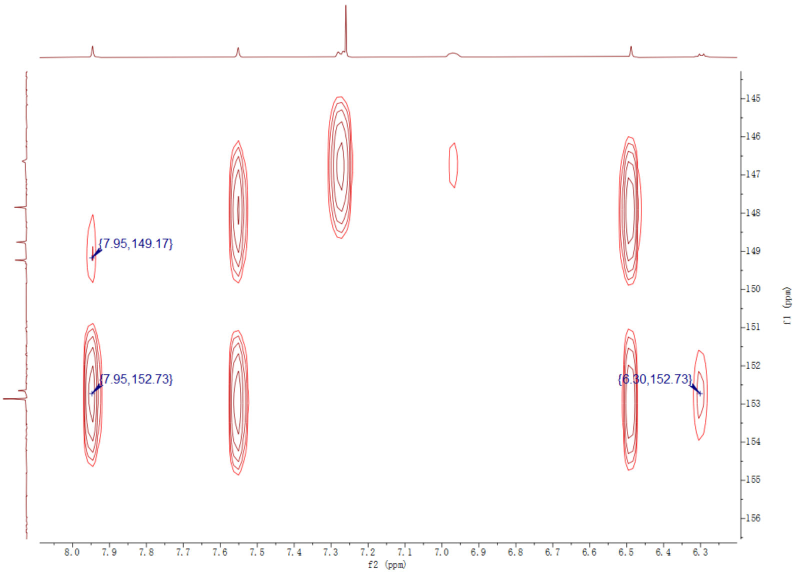


HMBC spectrum of **7**(CRH) in $CDCl_3$ zoom in

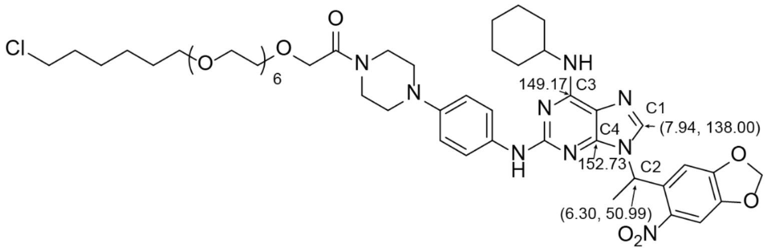


Structure determination rational: C1 has HMBC signal with C3 and C4. C4 is three bond distance and has stronger coupling signal compared to C3. Therefore, C3 is 149.17 ppm while C4 is 152.73 ppm. After looking into HMBC signal of C2, only C4 within three bonds distance from C2 shows up while C3 is five bonds away and does not show up.

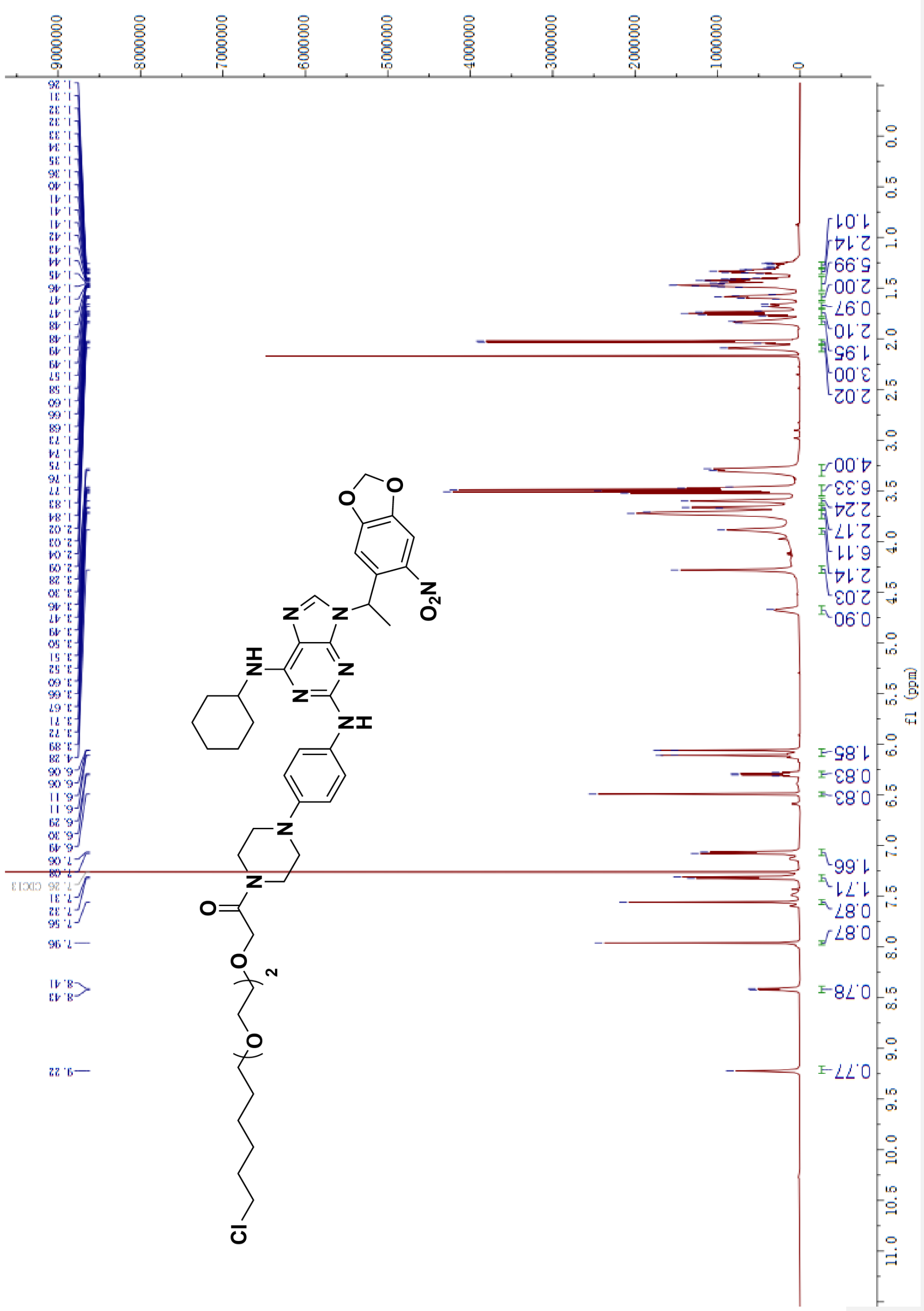

[1]H NMR spectrum of **8**(CRH-PEG2) in $CDCl_3$(600 MHz)

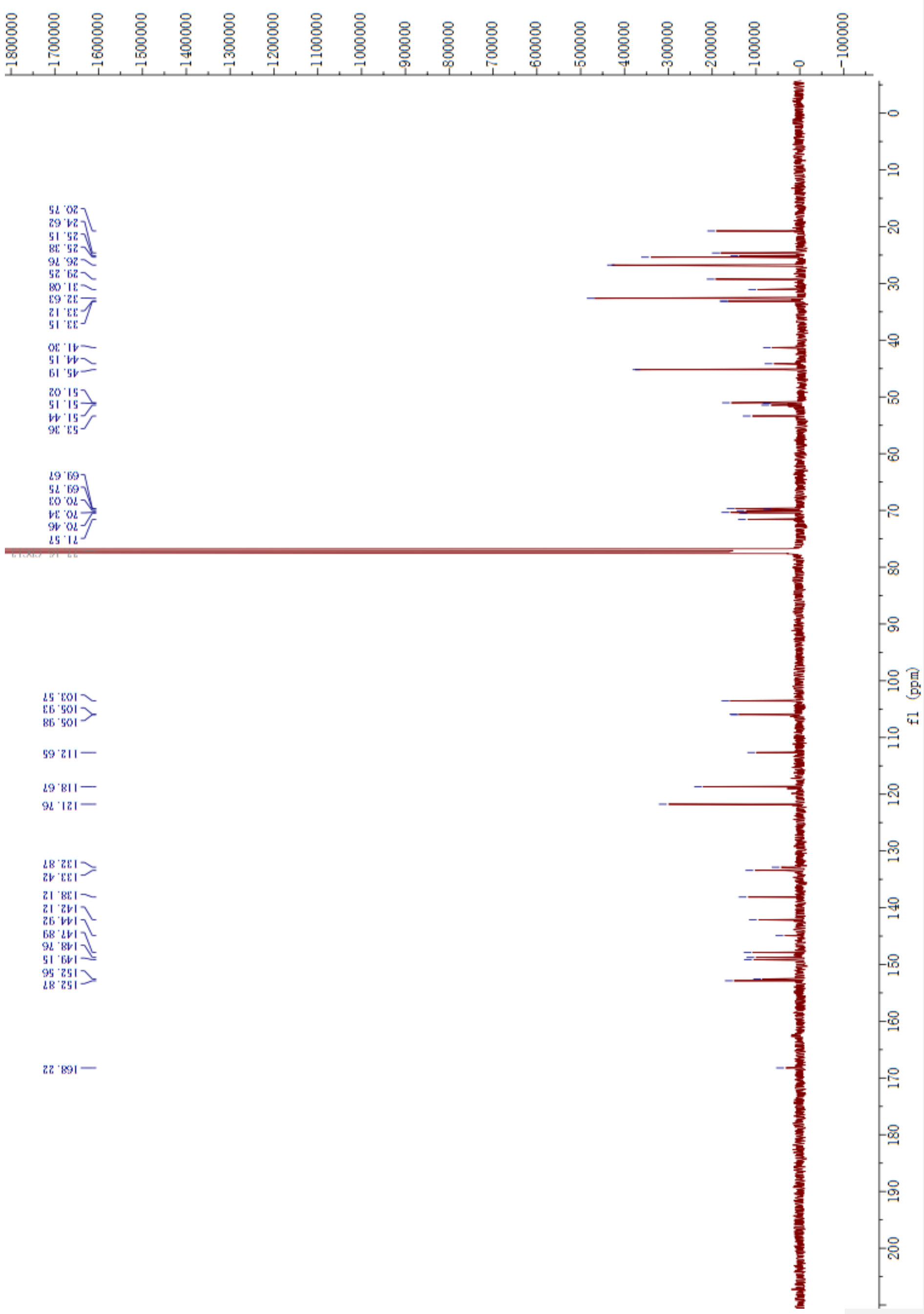


$^{13}C$ NMR spectrum of **8**(CRH-PEG2) in $CDCl_3$(151 MHz)

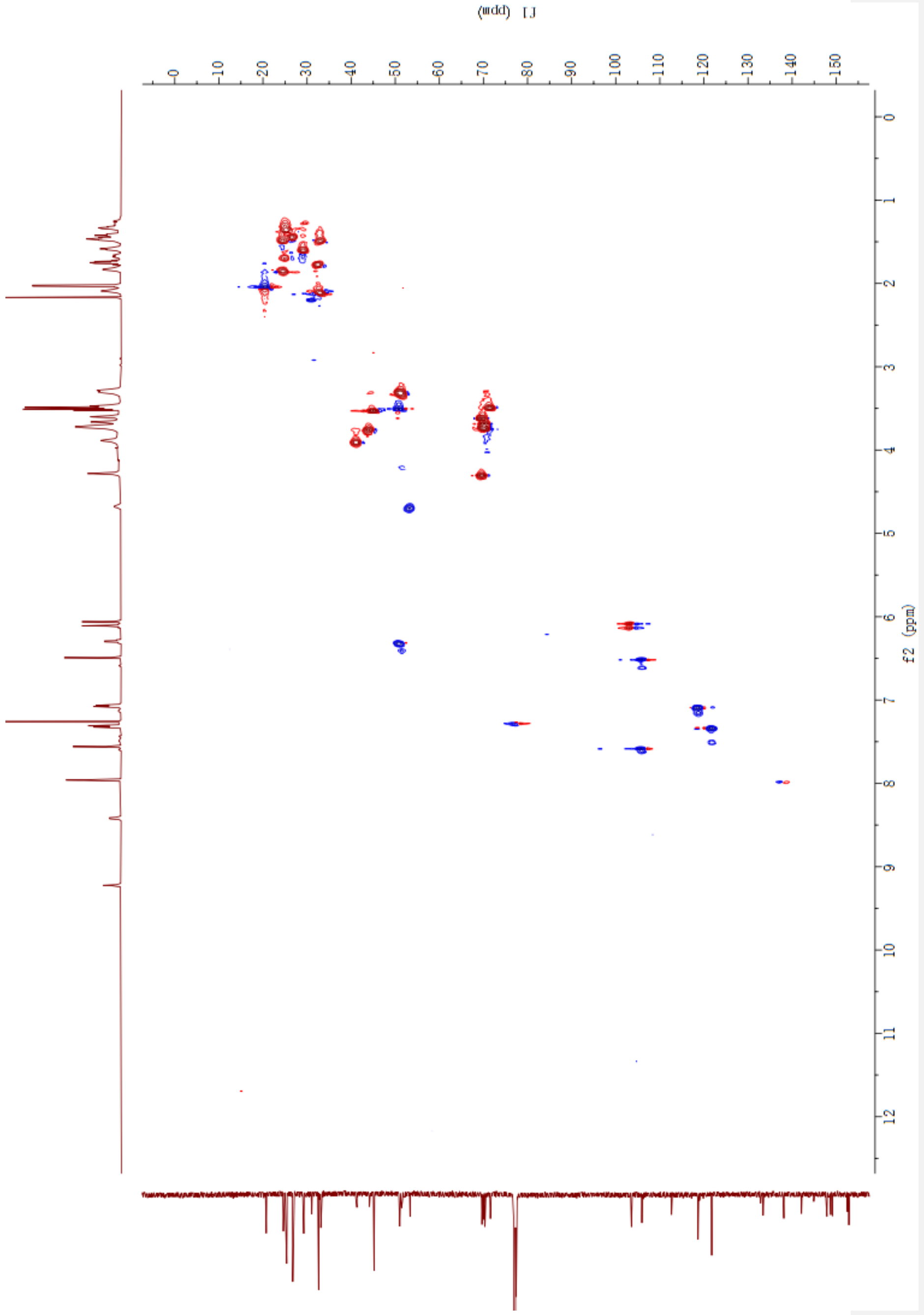


HSQC spectrum of **8**(CRH-PEG2) in $CDCl_3$

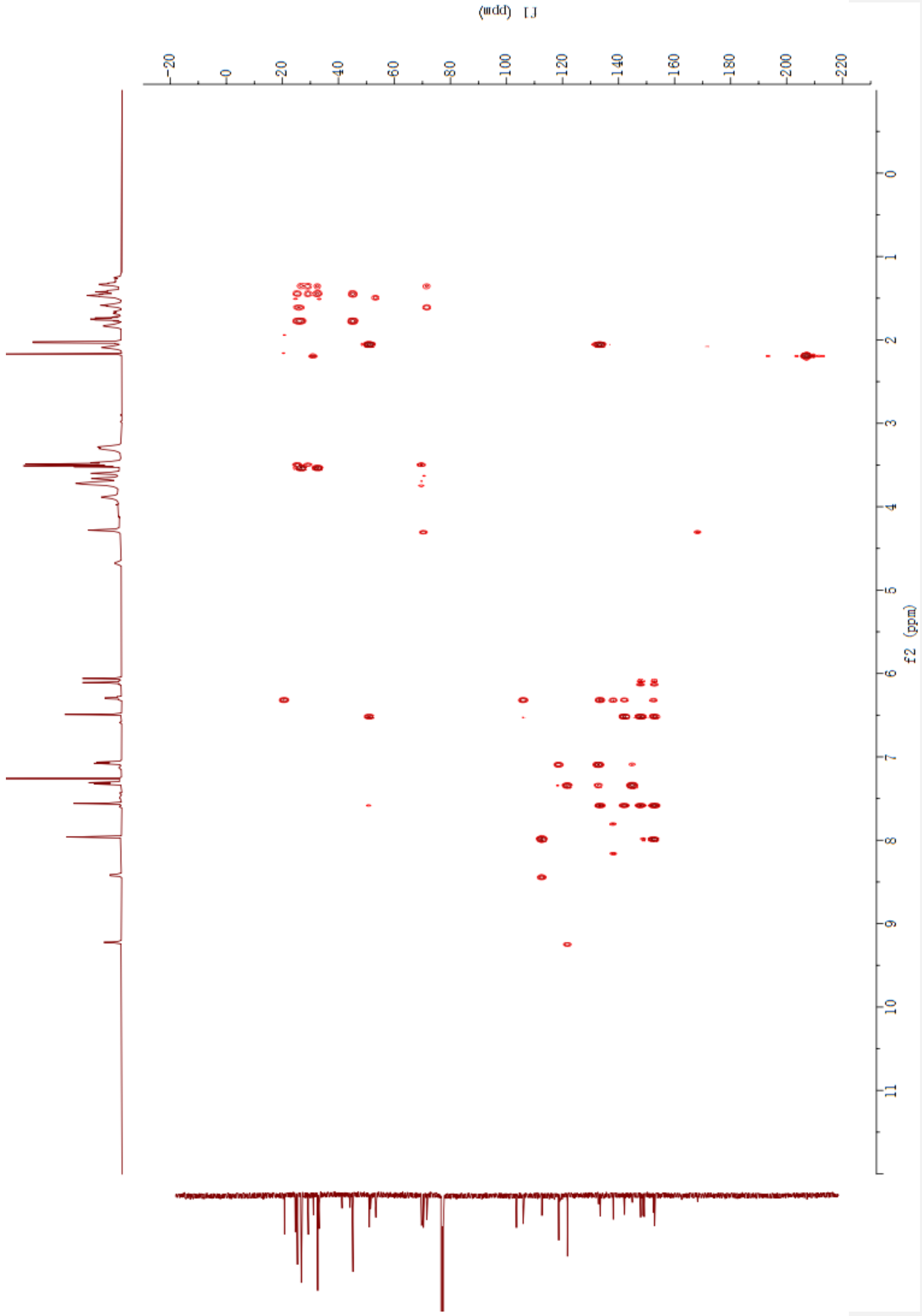


HMBC spectrum of **8**(CRH-PEG2) in $CDCl_3$

**Deleted:** 4
